# Long Noncoding RNA Derived from LncRNA–mRNA Co-expression Networks Modulates the Locust Phase Change

**DOI:** 10.1101/2020.04.11.036848

**Authors:** Ting Li, Bing Chen, Pengcheng Yang, Depin Wang, Baozhen Du, Le Kang

**Author notes:** Corresponding author. (Kang L).

## Abstract

Long noncoding RNAs (lncRNAs) regulate various biological processes from gene expression to animal behavior. Although protein-coding genes, microRNAs, and neuropeptides play important roles in the regulation of phenotypic plasticity in migratory locust, empirical studies on the function of lncRNAs in the process remain limited. Here, we applied high-throughput RNA-seq to characterize the expression patterns of lncRNAs and mRNAs in the time course of the locust phase change. LncRNAs displayed more rapid response at the early stages of the time-course treatments than mRNA expression. Functional annotations demonstrated that early changed lncRNAs employed different pathways in isolation and crowding processes to cope up with the changes in population density. Finally, two overlapping hub lncRNA loci in the crowding and isolation networks were screened to be functionally verified. Experimental validation indicated that LNC1010057 could act as a potential regulator to modulate the locust phase change. This work offers new insights into the mechanism underlying the locust phase change and expands the scope of lncRNA functions in animal behavior.

## Introduction

Long noncoding RNAs (lncRNAs) are the largest class of ncRNAs with length of longer than 200 nucleotides and lack coding potentials [1]. Similar to mRNAs, lncRNAs are transcribed by RNA polymerase II and can be capped, polyadenylated, and spliced. However, the averaged expression level and sequence conservation of lncRNAs are lower than those of mRNAs, resulting in the difficult detection and annotation of lncRNAs through conserved sequences [2,3]. With the development of high-throughput deep sequencing technology, thousands of lncRNAs were identified in many species and confirmed as crucial regulators for biological processes instead of byproducts of transcription [4–6]. In fact, the expression levels of protein-coding genes can be regulated by lncRNAs in a variety of biological processes, including mRNA transcription, stability, translation, and post-translational modification [7]. LncRNAs modulate the expression of protein-coding genes through *cis*-acting on neighboring genes and *trans*-acting on distal genes [8]. Thus, the expression of lncRNAs is correlated with the potential target genes and the function of lncRNAs can be predicted by the associated mRNAs in lncRNA–mRNA co-expression networks [9]. However, in different biological processes, the expression profiles of lncRNAs and mRNAs follow concordant or discordant patterns, suggesting the diverse interaction patterns between lncRNAs and mRNAs [10].

The lncRNA expressions display tissue specificity and are especially abundant in the nervous system [11], and increasing evidences suggest that lncRNAs play roles in nervous system development and function [12,13]. Several studies have demonstrated that lncRNAs can serve as vital regulators in modulating the behavior of mammals and insects. For example, upregulation of the lncRNA MEG3 improves spatial learning and memory capability of rats with Alzheimer’s disease through inactivating the PI3K/Akt signaling pathway [14]. The decline of lncRNA *Gomafu* induces anxiety-like behavior in mouse by regulating the expression of schizophrenia-related genes [15]. In *Drosophila*, the lncRNA *CRG* maintains the normal locomotor and climbing capabilities through regulating the transcription of the neighboring *CASK* gene, and another lncRNA *yar* regulates the night sleep time [16,17]. Nonetheless, the regulation of lncRNAs on animal behavior remains a fascinating and growing research field for scientific investigation.

Although less studied than in mammals and *Drosophila*, lncRNAs have been identified in non-model insect species [18,19]. In these studies, the expressions of lncRNAs were analyzed and functional annotation revealed that insect lncRNAs were involved in biological processes, such as insecticide resistance, fecundity, and gland apoptosis [19–21]. Particularly, four lncRNAs (i.e., *Ks-1, AncR-1, kakusei*, and *Nb-1*) in *Apis mellifera* are expressed preferentially in the brain and related to social behaviors [22]. However, no specific lncRNA has been experimentally confirmed to regulate behaviors in non-model insects.

The migratory locust, *Locusta migratoria*, is a worldwide agriculture pest and displays dramatic phenotypic plasticity, with the morphological, physiological, and behavioral traits reversely changing between the solitarious and gregarious phases [23]. Under high population density, gregarious locusts form large swarms, become attracted by conspecifics, exhibit active movement, and migrate long distances. By contrast, solitarious locusts live in an individual state, stay quiet, and are cryptically colored to blend with the surroundings. However, the two phases of locust have the entirely same genome which is 6.5G huge-genome with a very small portion of encoding proteins, and most of the genome regions express ncRNAs [24]. The molecular regulatory mechanism underlying two-phase changes include several protein-coding genes that are involved in the olfaction [25], dopamine biosynthesis and release pathway [26], and neuropeptide F/nitric oxide pathway [27,28]. Moreover, microRNAs also modulate phase-related traits by regulating the expressions of phase-related genes [29,30]. More recently, phase-, habitat-, and gender-specific lncRNAs were identified in the migratory locusts, implying the possible roles of lncRNAs in the locust phase change [31]. However, the lncRNA expression and function related to the locust phase change are still unknown.

To identify the phase-related lncRNAs, we systematically characterized the expression of locust lncRNAs and annotated their functions. In crowding and isolation processes, the expressions of lncRNAs were more temporally specific than those of mRNAs, and the response to phase change was more sensitive than that of mRNAs. Finally, a hub lncRNA derived from the co-expression between early changed lncRNAs and mRNAs was experimentally validated to modulate the phase-related behaviors. Our findings reveal the important roles of lncRNAs in the locust phase change and interactions between protein-coding genes and noncoding RNAs in the phenotypic plasticity of locusts.

## Results

### Identification and characterization of locust lncRNAs

To identify phase-related lncRNAs systematically, we constructed 24 rRNA-depleted RNA libraries with three biological replications for each treatment (**Figure 1A**). These samples were obtained from the locust brains undergoing time-course crowding and isolation treatments. In the crowding of solitarious locusts (CS), the solitarious locusts were kept in the same cage with gregarious locusts for 0, 4, 8, and 16 h to promote a behavioral transition from the solitarious toward the gregarious phase. In contrast to the CS process, the isolation of gregarious locusts (IG) was performed by individually keeping the gregarious locusts in separate cages for 0, 4, 8, and 16 h. The strand-specific RNA-seq on these libraries was performed on Illumina HiSeq 3000 platform.

**Figure 1.**
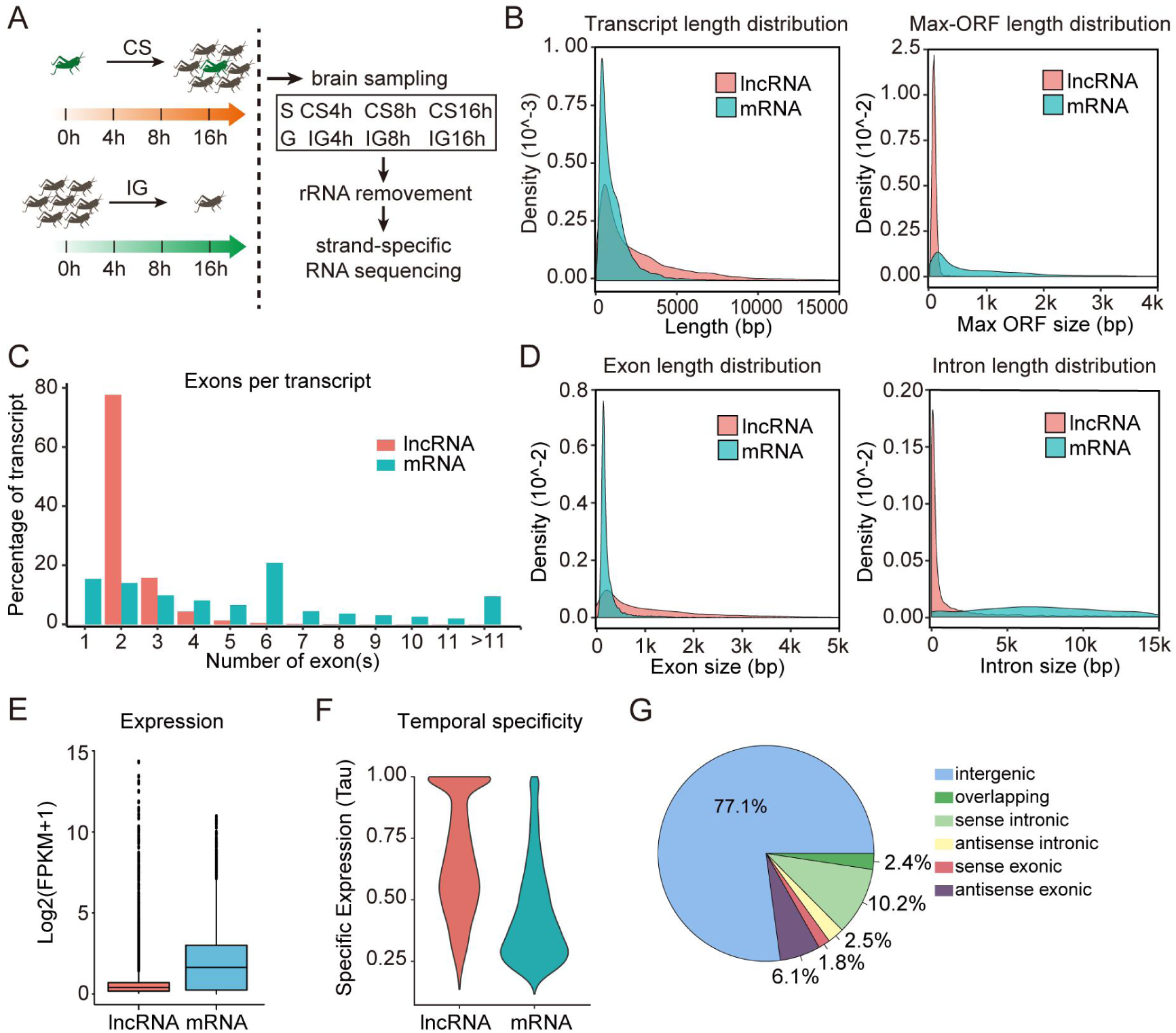
Identification and characterization of lncRNAs in the migratory locust. **A**. Experimental outline for the crowding and isolation treatments, sampling, and sequencing. G, gregarious locust; S, solitarious locust. CS, crowding of solitarious locusts; IG, isolation of gregarious locusts. The black vertical lines represent sampling time points. **B**. Full-length and Maximum open reading frame (ORF) size distribution of lncRNA and mRNA in migratory locusts. **C**. Number of exons per transcript for lncRNAs and mRNAs. **D**. Density distribution of exon size and intron size for lncRNAs and mRNAs. **E**. Overall expression (log_2_(FPKM+1)) of lncRNAs compared with mRNAs in the locust brain. **F**. Temporally specific expressions of lncRNAs and mRNAs in the time course. **G**. Locust lncRNA category and their proportions.

After removing low-quality reads, 363.14 Gb clean data of all samples were obtained, with the Q30 higher than 91.1% evaluated by FastQC [32], indicating that all the subsequent analyses were based on high-quality data (Table S1). A total of 1,597,688 transcripts from 1,431,906 loci were identified (Figure S1A). Sequencing saturation assessment indicated that the sequencing depth was sufficient to identify novel transcripts (Figure S1B). The expression levels of transcripts were measured as fragments per kilobase of transcript per million mapped reads (FPKM) by the RSEM method.

Based on the annotation of locust genome sequences [24], 15,309 known protein-coding transcripts were identified from the assembled transcripts using the Cuffcompare program in the Cufflinks package. The unknown transcripts were used for the putative lncRNA screening, and the analysis pipeline was developed to identify bona fide lncRNAs (Methods and Figure S1A). Finally, 14,373 highly reliable putative lncRNAs (FPKM > 1.0) from 10,304 loci were identified (Figure S1C and Table S2). The expression levels of lncRNAs were verified by quantitative reverse transcription-polymerase chain reaction (qRT-PCR) and displayed high correlation (Pearson’s *r* = 0.86) with those from RNA-seq (Figure S2).

We characterized the genomic features of locust lncRNAs by comparing them with the protein-coding mRNAs assembled in this study. The putative lncRNAs ranged from 202 bp to 25,251 bp in length, with a median length of 1,568 bp. The lncRNAs were significantly longer than mRNAs whose median length was 786 bp (Kolmogorov-Smirnov test [ks-test], D = 0.30002, *P* < 2.2e-16; **Figure 1B**, left). However, the lncRNAs are shorter than mRNAs in the size of the maximum open reading frame (ORF) predicted (median maximum ORF length of 78 bp for lncRNAs and 544 bp for mRNAs; ks-test, D = 0.71984, *P* < 2.2e-16; Figure 1B, right). For the number of exons contained in each transcript, approximately 78% of lncRNAs comprised 2 exons, whereas mRNAs contained exons ranging from 1 to 120 (**Figure 1C**). Thus, lncRNA exons are robustly longer than those of mRNA (average length: 1142 vs 263 bp; ks-test, D = 0.53967, *P* < 2.2e-16; **Figure 1D**, left). Meanwhile, lncRNA introns are significantly shorter than those of mRNA (average length: 10,086 vs 12,442 bp; ks-test, D = 0.61789, *P* < 2.2e-16; Figure 1D, right). The expression level analysis indicated that the global expression of lncRNAs was significantly lower than that of mRNAs (average of 0.6 vs 1.9; Student’s t-test, *t* = 79.868, *P* < 2.2e-16; **Figure 1E**). However, the temporal specificity analysis demonstrated the higher specific expression for lncRNAs than mRNAs (average of 0.672 vs 0.436; Student’s *t*-test, *t* = 79.868, *P* < 2.2e-16; **Figure 1F**). On the basis of lncRNA genomic locations relative to mRNAs, the locust lncRNAs are classified as intergenic, overlapping, sense intronic, antisense intronic, sense exonic, and antisense exonic. More than 77% of locust lncRNAs were long intergenic noncoding RNA (lincRNA, **Figure 1G**). Collectively, these results indicate that the locust lncRNAs differ considerably from its mRNAs in terms of structure and expression. The locust lncRNAs were longer, possessed less but longer exons and shorter introns. Their expressions displayed higher temporal-specificity than mRNAs.

### Higher specific expression of lncRNAs in gregarious locusts compared to solitarious locusts

A total of 9722 lncRNAs (73.9%) were commonly expressed in both solitarious and gregarious locust brains, whereas 962 lncRNAs (7.3%) were expressed specifically in solitarious locusts and 2469 (18.8%) in the gregarious (**Figure 2A**, top). However, only 479 and 1090 mRNAs (3.1% and 7.1%) were expressed specifically in solitarious and gregarious locusts, respectively (Figure 2A, bottom). These results indicate that higher percentage of lncRNAs were expressed specifically in the two locust phases compared with mRNAs, and more lncRNAs were expressed in gregarious than in solitarious locusts. Compared with the expressions in solitarious locusts, 335 lncRNAs and 779 mRNAs were downregulated, whereas 313 lncRNAs and 261 mRNAs were upregulated in gregarious locusts (fold change > 2 and *P* < 0.05; **Figure 2B**). LincRNAs were the most regulated lncRNAs (81.2% and 87.8%) in down- and upregulated lncRNAs. Antisense exonic and sense intronic and antisense exonic lncRNAs were the second most down- and upregulated lncRNAs, respectively (Figure 2B). These lncRNAs and their nearby protein-coding genes were showed in Supplement Table 3, and the top 10 lncRNA genes ranked by fold change were displayed in **Figure 2C**. These results indicate that the number and the expressions of lncRNAs in solitarious and gregarious locust brains significantly differed.

**Figure 2.**
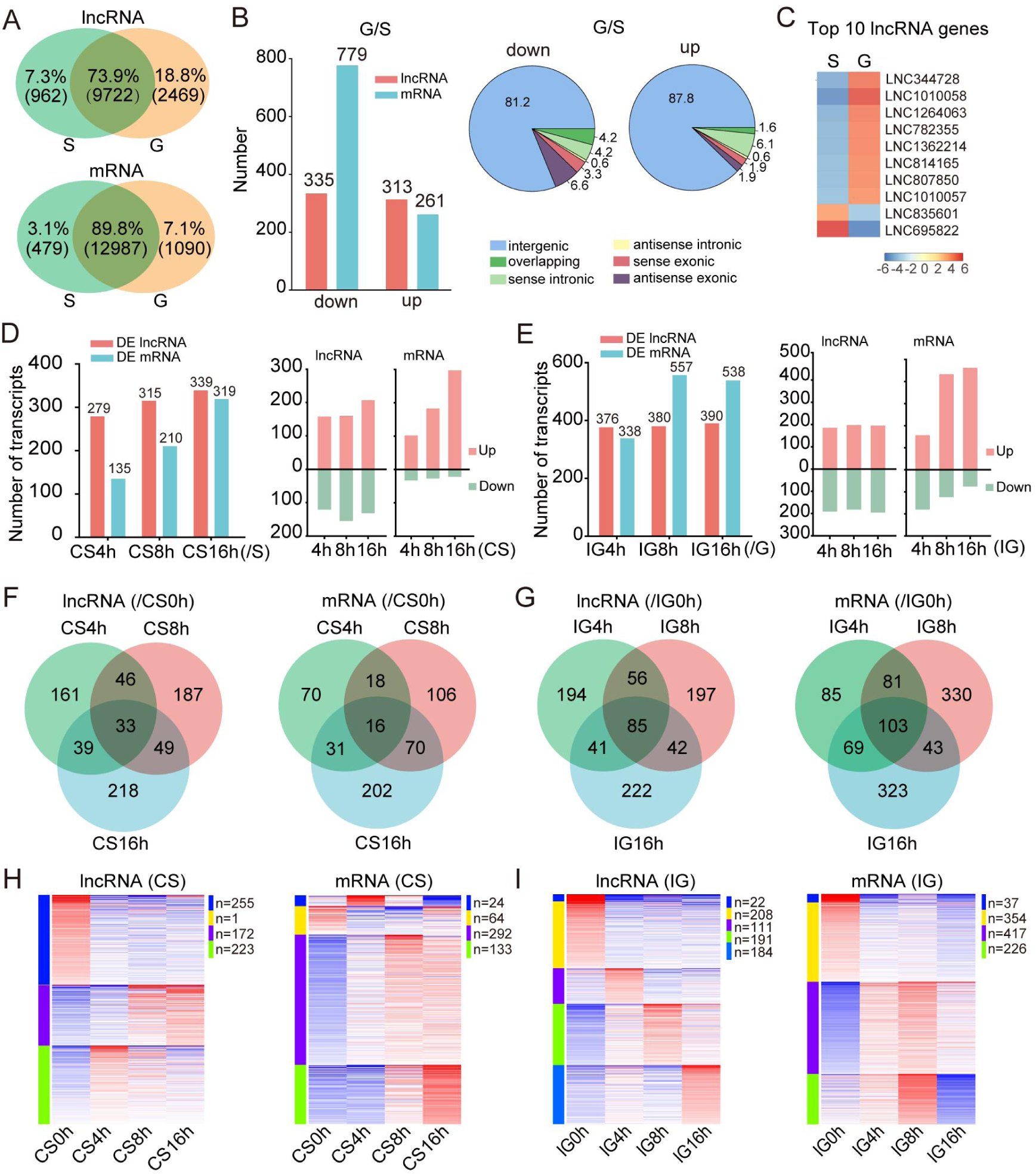
DE lncRNAs and mRNAs in CS and IG. **A**. Venn diagram of expressed lncRNAs and mRNAs in gregarious and solitarious locusts. **B**. Down- and upregulated lncRNAs and mRNAs in gregarious locusts compared with those in solitarious locusts. Classification of DE lncRNAs between G and S was on the right. **C**. The top 10 DE lncRNAs between gregarious and solitarious locusts. **D**. The number of total regulated, up- and downregulated lncRNAs and mRNAs at each time point relative to 0 h in CS. **E**. The number of total regulated, up- and downregulated lncRNAs and mRNAs at each time point relative to 0 h in IG. Venn diagram of DE lncRNAs and mRNAs in CS (**F**) and IG (**G**). K-mean clustering of DE lncRNAs and mRNAs in CS (**H**) and IG (**I**).

### LncRNAs as different signatures of phase change from mRNAs

At different points of the time-course samples of CS, the percentages of totally expressed lncRNAs ranged from 64.3% to 68.9%, whereas those for mRNAs ranged from 87.9% to 89.7% (Figure S3A, top left). During IG, the percentages of totally expressed lncRNAs ranged from 64.8% to 81.0%, whereas those for mRNA ranged from 87.2% to 91.9% (Figure S3A, top right). However, the proportions of specifically expressed lncRNAs in the same sample were higher than those of mRNAs during CS and IG (Figure S3A, bottom). The specific expression pattern of lncRNAs indicates its relationship with the locust phase change.

The differentially expressed (DE) lncRNAs and mRNAs during CS and IG were identified by comparing the normalized expression of transcripts at each point of time course with that at the 0 h time-point (i.e., CS4h/CS0h, CS8h/CS0h, CS16/CS0h, IG4h/IG0h, IG8h/IG0h, and IG16h/IG0h). A total of 1246 and 1871 transcripts were differentially expressed in CS and IG, respectively. Among these transcripts, 733 lncRNAs and 513 mRNAs changed expression in CS, whereas 837 lncRNAs and 1034 mRNAs changed expressions in IG (Figure S3B). These results indicate that more lncRNAs are involved in CS, whereas more mRNAs are engaged in IG. The DE lncRNAs and mRNAs at CS16h compared with G, and IG16h compared with S were also identified (Figure S3C). In this result, more mRNAs were involved in CS16/G and more lncRNAs were involved in IG16h/S. This result is contrary with the previous one but exactly indicate that the crowding and isolation treatments are effective. At each time point of CS and IG, the numbers of DE lncRNAs were little different but that of mRNAs were quite diverse (**Figure 2D**, left and **2E**, left). Moreover, the up- and downregulated lncRNAs at each point of time course in CS and IG displayed comparative numbers but the upregulated mRNAs were considerably more than the downregulated mRNAs in CS and IG (Figure 2D, right and 2E, right). Identically, the numbers of constantly changed transcripts in the time course were different for lncRNAs and mRNAs. More lncRNAs were constantly changed than mRNAs in CS but less than mRNAs in IG (lncRNAs vs mRNA: 33 vs 16 in CS and 85 vs 103 in IG; **Figure 2F and 2G**). Additionally, the K-means clustering of the lncRNA expression differed with that of mRNAs during both CS and IG (**Figure 2H and 2I**). All the data suggest that lncRNAs respond differently to both the crowding and isolation treatments in comparison with mRNAs. However, hierarchical clustering analyses demonstrated that DE lncRNAs and mRNAs clustered the same time-point samples together in CS and IG, suggesting that both lncRNA and mRNA can be signatures of the locust phase change (Figure S4).

### Rapid response of lncRNAs to the population density change

With the time course of the crowding or isolation treatments, the number of DE lncRNAs and mRNAs increased (Figure 2D and 2E). However, the percentage of the regulated lncRNAs was 2/5 higher than that of mRNAs at the 4 h time-point in CS (Chi-Square test, *P* = 0.002) and 1/3 higher than that of mRNA at the 4 h time-point in IG (Chi-Square test, *P* = 3.62e-4; **Figure 3A**). Meanwhile, the number of changed lncRNAs reached the highest level at a much earlier stage (4 h) than mRNAs (8 h or later). Identically, the percentage of specifically changed lncRNAs at the 4 h time-point was also higher than that of mRNAs in CS and IG (Chi-Square test, *P* = 0.008 in CS and *P* = 3.28e-13 in IG) and reached a relative stable stage more rapidly (**Figure 3B**). Similar results were observed in the upregulated lncRNAs and mRNAs in IG (Chi-Square test, *P* = 0.002) and downregulated lncRNAs in CS and IG (Chi-Square test, *P* = 3.30e-6 in CS and *P* = 0.02 in IG; **Figure 3C**). These results indicate that lncRNAs respond more rapidly to crowding and isolation treatments than mRNAs.

**Figure 3.**
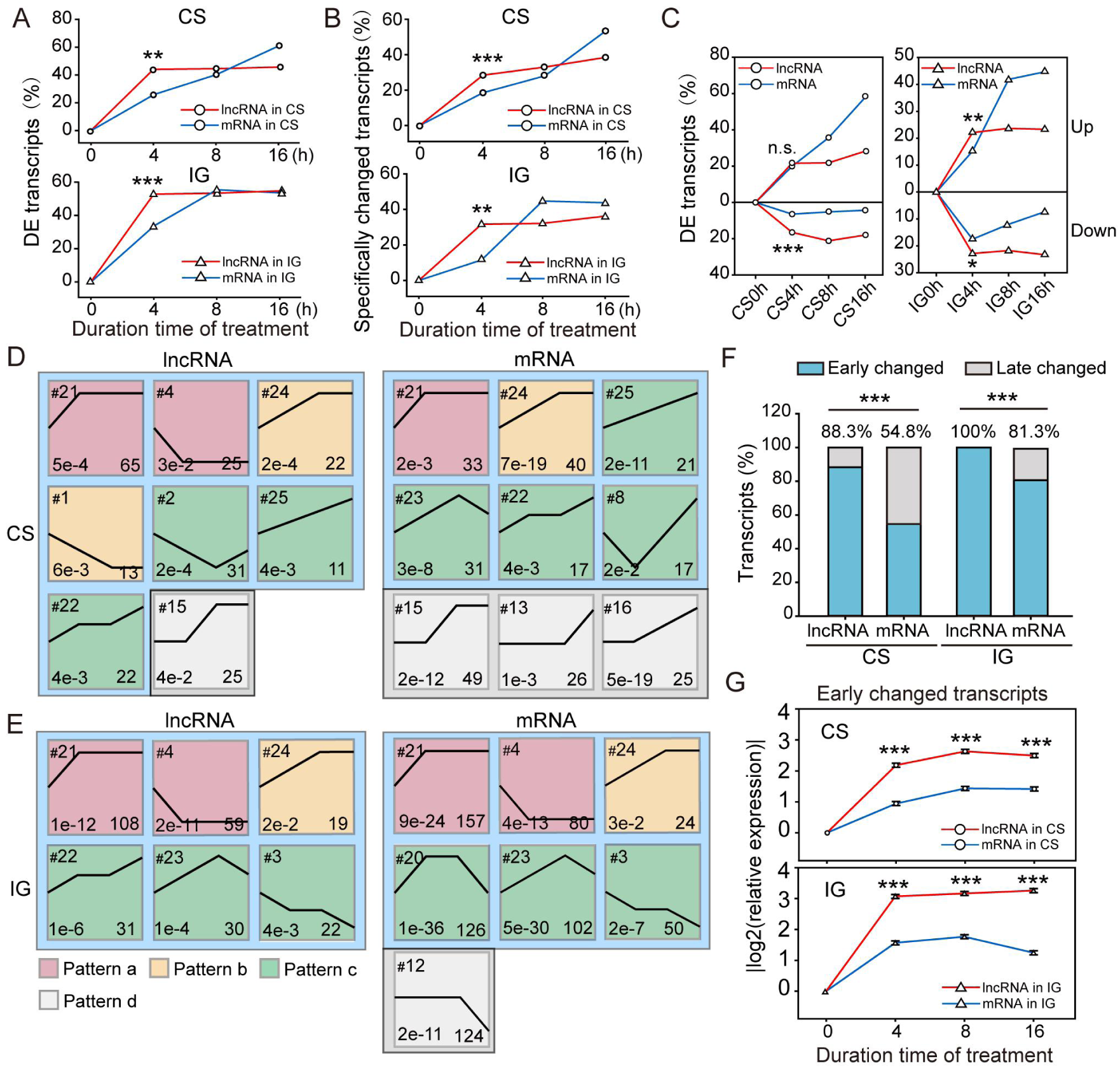
Different responses of lncRNAs and mRNAs to crowding and isolation treatments. **A**. Percentage of the DE lncRNAs and mRNAs at each time point of time course relative to total DE lncRNAs and mRNAs in CS and IG. **B**. Percentages of specifically changed lncRNAs and mRNAs at each time point of time course relative to total specifically expressed lncRNAs and mRNAs in CS and IG. **C**. Percentages of up- and downregulated lncRNAs and mRNAs at each time point of time course in CS (left) and IG (right) processes relative to the total DE lncRNAs and mRNAs. Significant expression profiles (*P* < 0.05) of lncRNAs and mRNAs clustered via STEM software in CS (**D**) and IG (**E**). The numbers in the left upper part of boxes are profile serial numbers, those in the left lower part are p-values, and those in the right lower part are numbers of transcripts contained in profiles. Pattern a (early-changed) profiles are violet, Pattern b (early-middle-changed) profiles are yellow, Pattern c (sustainably-changed) profiles are green, and Pattern d (late-changed) profiles are gray. **F**. Percentages of the early and late changed transcripts (lncRNAs or mRNAs) of CS and IG. **G**. Expression patterns of early changed lncRNAs and mRNAs during CS and IG. The expressions levels are relative expressions (|log2(v(*i*)/v(0))|) of all early changed lncRNAs or mRNAs. V(*i*) and v(0) represent the expressions of the transcripts at *i* h (*i* = 4, 8, 16) time point and 0 h time point, respectively. Significant differences between lncRNAs and mRNAs in **A, B, C**, and **F** were measured through Chi-squared test. Data in G are shown as mean ± SE. \**P* < 0.05, \*\**P* < 0.01, \*\*\**P <* 0.001.

To further analyze the expression patterns of lncRNAs and mRNAs, we used Short Time-series Expression Miner (STEM) to cluster transcripts on the basis of their expressions. In STEM analysis, twenty-six model profiles were set to describe the expression patterns of the regulated lncRNAs and mRNAs in CS and IG, but only several were screened as significant expression profiles (*P* < 0.05, Figure S5). During CS, 214 lncRNAs and 242 mRNAs were clustered into eight and nine significant expression profiles, respectively (**Figure 3D**). Among these profiles, lncRNA expression profile 15 and mRNA expression profiles 15, 16, and 13 displayed no expression changes until 8 or 16 h after the crowding treatment. Thus, we termed them as late changed profiles (Pattern d). In contrast to late changed profiles, early changed profiles featured transcript expression that started changing at the 4 h time-point. In accordance with the expression change after the 4 h time point, the early changed profiles were subdivided into Pattern a (early-changed), Pattern b (early-middle-changed), and Pattern c (sustainably-changed). During the IG process, 269 lncRNAs and 639 mRNAs were clustered into six and seven significant expression profiles, respectively, in which all lncRNA profiles were the early changed (**Figure 3E**). Among the mRNA expression profiles, four were the early changed, but profile 12 was the late changed profile. The numbers of clustered profiles for lncRNAs showed no significant differences with those for mRNAs in CS and IG. However, by calculating the transcripts number in profiles, we found that the percentage of early changed lncRNAs was 61.1% higher than that of mRNA in CS (88.3% vs 54.8%; Chi-Square test, *P* = 1.05e-7; **Figure 3F**). Similarly, the percentage of early changed lncRNAs was also higher than that of mRNAs in IG (100% vs 81.3%; Chi-Square test, *P* = 3.93e-12; Figure 3F). Thus, lncRNAs responded sooner to the crowding and isolation treatments than mRNAs in general.

Additionally, the expression change rate of the early changed lncRNAs was faster than that of mRNAs in CS (0.55 vs 0.24; Student’s *t*-test, *t* = 7.88, *P* = 9.32e-14) and IG (0.77 vs 0.39; Student’s *t*-test, *t* = 10.42, *P* = 2.32e-22; **Figure 3G**). Therefore, both the ratio and the expression change rate of early changed lncRNAs were much higher than that of early changed mRNAs, indicating that locust lncRNAs were more sensitive to the changes in the population density than mRNAs.

### LncRNAs involved in different pathways in the early-period changes of CS and IG

To determine the functions of the early changed lncRNAs and hub lncRNAs in the locust phase change, we analyzed the correlations between lncRNAs and mRNAs by constructing co-expression networks. The Pearson’s correlation coefficient between early changed lncRNAs and all detectable mRNAs was evaluated in CS and IG. Only the lncRNA–mRNA pairs with strong correlation (|*r*| > 0.9) were used to construct the networks (**Figure 4A** and **4B**, Table S4). In the co-expression networks, the nodes and correlations were divided into three modules, that is, Pattern a, Pattern b, and Pattern c module, on the basis of the lncRNA expression patterns defined above. Coupled with lncRNAs, correlated protein-coding genes were separated into relevant modules. The Kyoto Encyclopedia of Genes and Genomes (KEGG) enrichment of the mRNAs in each module was analyzed to predict the biological pathways in which the lncRNAs might be involved.

**Figure 4.**
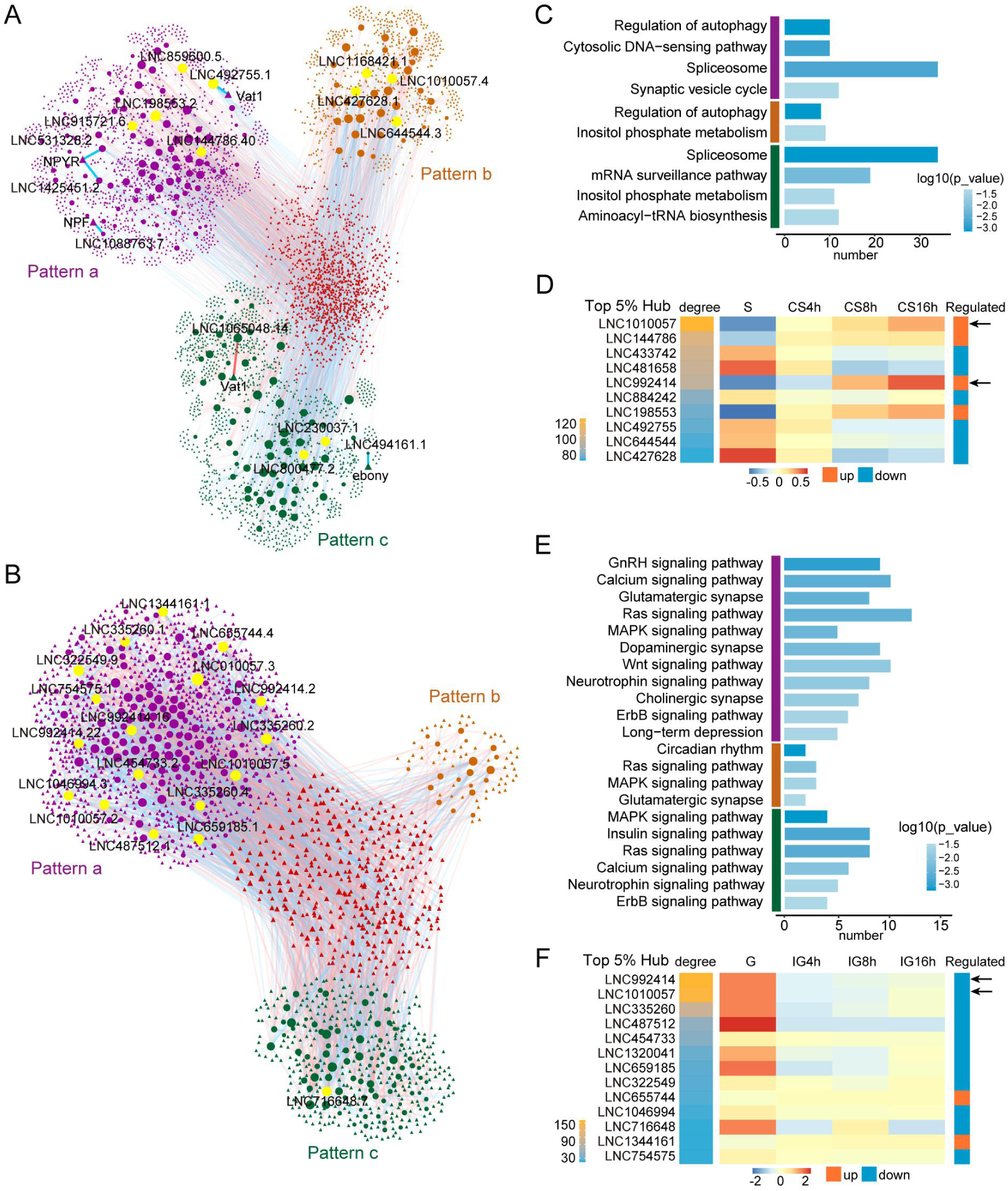
Functional prediction and hub lncRNA identification. Co-expression networks between lncRNA and protein-coding genes constructed in CS (**A**) and IG (**B**) by using Pearson correlation coefficient. The circular nodes and squares represent lncRNAs and mRNAs, respectively. The red and blue lines denote positive and negative correlations, respectively. The nodes in the network were divided into three modules, including Pattern a (violet), Pattern b (yellow), and Pattern c (green) modules, based on the lncRNA expression patterns. The lncRNAs with top-ranked degrees are colored yellow. **C**. KEGG pathways were enriched for each module in the CS network. Colored vertical bars represent the different expression modules. **D**. Top 5% of hub lncRNA loci ranked by degree in CS. The heatmap represents the expression changes of hub lncRNA loci in CS. **E**. KEGG pathways were enriched for each module in the IG network. **F**. Top 5% of hub lncRNA loci ranked by degree in IG. The heatmap represents the expression changes of hub lncRNA loci in IG. The black arrows indicate the overlapping hub lncRNAs in CS and IG networks.

In CS, “Regulation of autophagy”, “Spliceosome”, and “Inositol phosphate metabolism” pathways were overrepresented in the top rank and in at least two modules (**Figure 4C**). The results indicate that early changed lncRNAs in CS play important roles in regulating autophagy, RNA splicing, and signal transduction pathways. The “Regulation of autophagy” pathway was involved in the initial stage for early-changed and early-middle-changed lncRNAs, whereas those for “Spliceosome” and “Inositol phosphate metabolism” were both involved for sustainably-changed lncRNAs. Of note, “Synaptic vesicle cycle” pathway, which is related to neurotransmitter release, was only overrepresented for lncRNAs in Pattern a (early-changed) module (Figure 4C). This result implies that early-changed lncRNAs may have the more important roles in regulating the nervous system during CS. In addition, several lncRNAs were strongly associated with known phase-related genes. For instance, early-changed lncRNAs LNC531328.2, LNC1425451.2, LNC1088763.7, and LNC492755.1 were associated with *NPYR, NPF1a*, and *Vat1* genes [28] (*r* = -0.90, -0.91, -0.98, -0.95; Figure 4A, Table S5). The sustainably-changed lncRNAs LNC494116.1 and LNC1065048.14 were associated with *Ebony* and *Vat1* genes in the dopamine pathway [26] (*r* = -0.92, 0.98).

In the lncRNA–mRNA networks, a lncRNA is more important if the degree value describing the connection number is higher. In the CS network, the top 5% lncRNAs in terms of the degree value were regarded as hub lncRNAs (Figure 4A). At the same time, the degree values of lncRNA loci were calculated and ranked to identify hub lncRNAs in “gene” level because several lncRNAs are derived from the same lncRNA locus, and lncRNA genes can be functional units [33]. In CS, 10 lncRNA loci that were ranked in the top 5% degree values were identified as hub lncRNA genes, of which four were upregulated and the other six were downregulated (**Figure 4D**).

During IG, the synapse related pathways, including “Glutamatergic synapse”, “Dopaminergic synapse”, and “Cholinergic synapse”, were enriched. Most synapse related pathways were enriched in Pattern a (early-changed) module. Meanwhile, the “Dopaminergic synapse” and “Cholinergic synapse” pathways were only involved in Pattern a (early-changed) module (**Figure 4E**). Signal transduction pathways, such as “Calcium signaling pathway”, “Ras signaling pathway”, “Insulin signaling pathway”, and “MAPK signaling pathway” were also enriched in IG (Figure 4E). The results of functional annotations suggest that early changed lncRNAs participate more in synapse related and signal processing pathways in IG than in CS.

The top 5% lncRNAs in the degree value were regarded as hub lncRNAs in the IG network (Figure 4B), and the number of hub lncRNAs in the Pattern a (early-changed) module was the highest, suggesting that the early-changed lncRNA play important roles in the locust phase change. At the same time, 13 hub lncRNA loci that ranked in the top 5% degree values were identified in IG (**Figure 4F**). Among these hub lncRNA loci, two were upregulated and the other eleven were downregulated. Interestingly, two overlaps were observed in the hub lncRNA loci identified in both CS and IG (Figure 4D and 4F). Moreover, their expressions increased in CS and decreased in IG, indicating that they sensitively responded to crowding and isolation treatments with different directions. Thus, LNC1010057 and LNC992414 were regarded as the valuable candidate regulators in the locust phase change.

### LNC1010057 was verified to regulate the locust phase change

To verify the function of LNC1010057 and LNC992414 in the locust phase change, we performed a series of molecular biology experiments. First, the longest transcripts identified in LNC1010057 and LNC992414 loci were cloned. LNC1010057 consisted of repeat A, B, and C elements, which were sequentially distributed and repeated four times, except the C element, which was repeated three times (**Figure 5A**). Intriguingly, sequence alignments demonstrated that LNC992414 shared similar repeat sequences with LNC1010057 in the 5′-end, but they were transcribed from different genome loci (Figure 5A and Figure S6). Despite the quantitative expression level of LNC992414 can be detected by specific primers, the expression level of LNC1010057 actually represent the total expression level of repetitive elements. As expected, the expression patterns of LNC1010057 and LNC992414 were extremely similar. During IG, LNC992414 and LNC1010057 significantly decreased after 4 h of isolation and remained relatively stable afterward (one-way ANOVA, F = 33.47, *P* = 5.45e-8; **Figure 5B**). By contrast, both lncRNAs displayed a consistently increasing trend and achieved a nearly 4-fold increase at 16 h time-point (one-way ANOVA, F = 5.22, *P* = 0.009; Figure 5B). The similarity of the sequence structure and expression pattern of LNC1010057 and LNC992414 suggest that they are homologous lncRNAs. However, the absolute real-time qPCR demonstrated that the LNC1010057 expression level was approximately 1,000-fold higher than that of LNC992414 in the brains of gregarious and solitarious locusts (**Figure 5C**).

**Figure 5.**
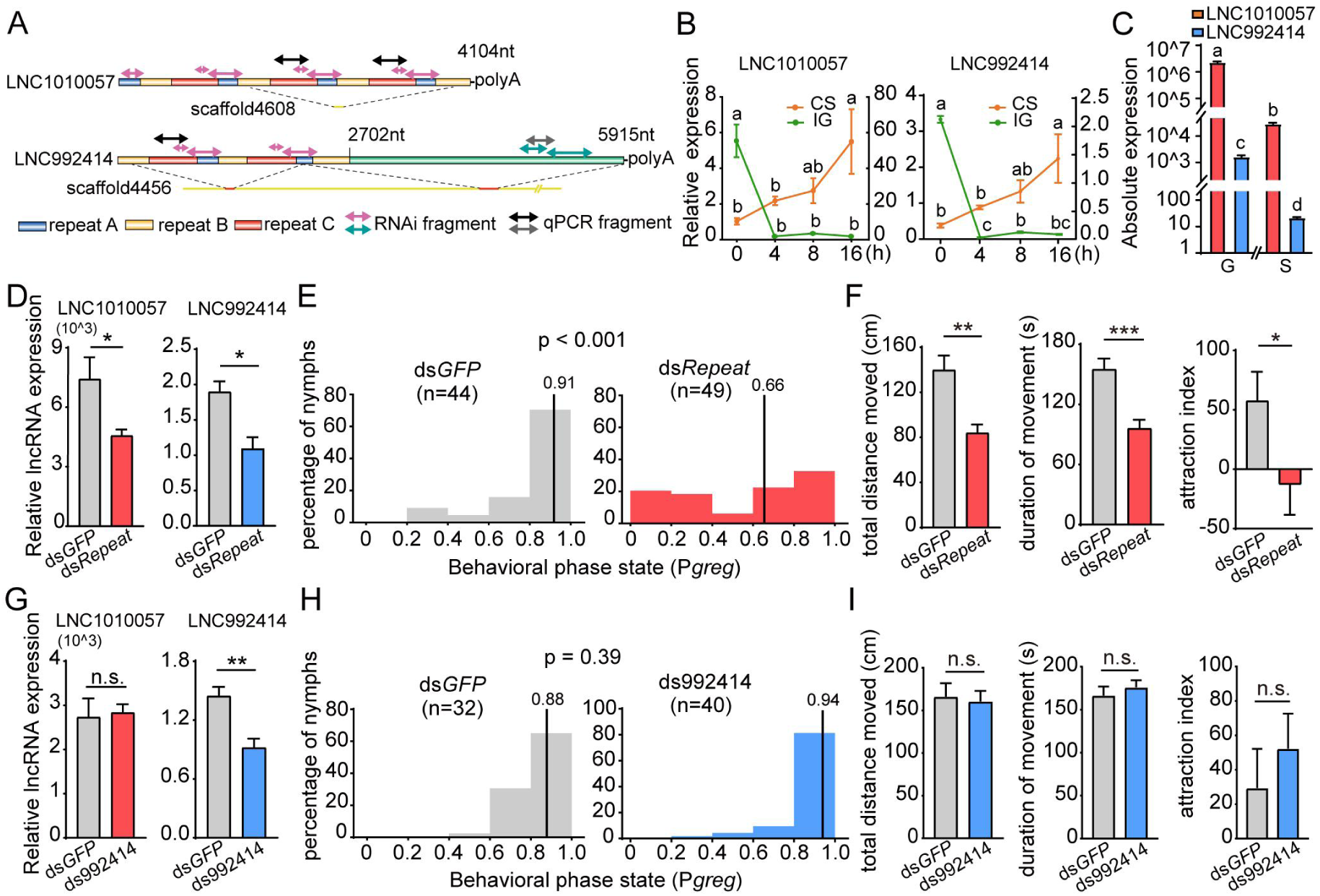
LNC1010057 potentially regulates the phase change of gregarious locusts. **A**. Full-length sequence structure and genome location of LNC1010057 and LNC992414. **B**. Expression changes of LNC1010057 and LNC992414 in the time course of CS and IG. Left and right coordinate axes show the relative expression in CS and IG, respectively. **C**. Absolute expression levels of LNC1010057 and LNC992414 in gregarious and solitarious locusts. **D**. Efficiency of the RNAi of LNC1010057 and LNC992414 in the locust brain after dsRNA injections. **E**. Phase changes in gregarious locusts when LNC1010057 and LNC992414 were knockdown in the brain. P_*greg*_, probabilistic metric of gregariousness. Vertical lines indicate median P_*greg*_ values. **F**. Changes in total distance moved, duration of movement, and attraction index of gregarious locusts after the knockdown of LNC1010057 and LNC992414. **G**. Efficiency of LNC992414 RNAi in locust brains after dsRNA injection. **H**. Phase changes in gregarious locusts when LNC992414 was knockdown in the brain. **I**. Changes in total distance moved, duration of movement, and attraction index of gregarious locusts after LNC992414 knockdown. Measurements are shown as mean ± SE. \**P <* 0.05, \*\**P <* 0.01, \*\*\**P <* 0.001 (Student’s *t*-test).

Second, to test whether LNC1010057 and LNC992414 functioned in the locust phase change, the behavioral assay was conducted after suppressing their expressions in locust brains by RNA interference (RNAi). The injection of double-strand RNA (dsRNA) to target the repetitive elements into gregarious brains significantly decreased the LNC1010057 (7470 vs 4610; Student’s *t*-test, t = 2.93, *P* = 0.02) and LNC992414 expression levels (1.9 vs 1.0; Student’s *t*-test, t = 3.88, *P* = 0.004; **Figure 5D**). The behavioral assay results demonstrate that the gregarious locusts significantly change their behaviors toward the solitarious state with the expression decrease of LNC1010057 and LNC992414 (Mann–Whitney U test, U = 599, *P* < 0.001, ds*Repeat* vs ds*GFP*; **Figure 5E**). Furthermore, multiple phase-related behavior parameters changed after the injection of repetitive element dsRNA. The total distance moved (TDM) and total duration of movement (TDMV) significantly decreased (Student’s *t*-test, t = 4.12, *P* < 0.001; t = 4.64, *P* < 0.001; **Figure 5F**). However, the movement speed showed no difference with the reduced repetitive element expression (Student’s *t*-test, t = 0.008, *P* = 0.93; Figure S7A). The conspecific preference behavior of gregarious locusts significantly decreased when the repetitive elements were knocked down (58.3 vs -13.5; Student’s *t*-test, t = 2.07, *P* = 0.04; Figure 5F). The main reason is that locusts spend shorter duration in the attraction zone (Student’s *t*-test, t = 2.34, *P* = 0.02) rather than longer duration in the repulsion zone (Student’s t-test, *t* = 1.14, *P* = 0.26; Figure S7B). In addition, the movement duration in the border zone significantly decreased (Student’s *t*-test, t = 3.44, *P* < 0.001). However, the difference in the central zone was insignificant (Student’s *t*-test, t = 1.76, *P* = 0.08) after the repetitive element expression decreased (Figure S7C). The turn times decreased (Student’s *t*-test, t = 3.06, *P* = 0.003), and the mean turn angle increased (Student’s *t*-test, t = 2.96, *P* = 0.004) compared with the those of the *GFP* control groups (Figure S7D). However, the specific knockdown of LNC992414 expression cannot cause the shift from the gregarious phase to the solitarious state (**Figure 5G** and **5H**). The phase-related behavior parameters, including the movement and the preference behavior for the conspecifics, showed no difference compared with the ds*GFP* control (**Figure 5I**, Figure S7E–S7H). Because the RNAi experiments for only LNC992414 did not cause the behavioral changes, the role of LNC992414 in regulating the locust phase change was excluded. However, no LNC1010057-specific RNAi is performed because the sequence of LNC1010057 is almost identical to the 5′-end of LNC992414. These results prove that LNC1010057 but not LNC992414 potentially regulates the locust phase change.

## Discussion

Here, we provided a time series lncRNAs expression profiling. Thus, we could identify the DE lncRNAs between gregarious and solitarious phases as well as during the locust phase change. We found that lncRNAs responded sooner to the change in population density than mRNAs. Base on the lncRNA–mRNA co-expression networks, the functions of the early changed lncRNAs were predicted and hub lncRNAs were selected. Finally, one hub lncRNA was experimentally confirmed to regulate the locust phase change.

### Features of locust lncRNAs

Locust lncRNAs possess different features compared with other insect species. One of the most significant characteristics is the longer transcript length of locust lncRNAs than mRNAs. In contrast, in other insect species, such as *Nilaparvata lugens* [20], *Apis mellifera* [18], and *Plutella xylostella*, the average transcript length of lncRNAs is shorter than that of mRNAs [19]. This result possibly results from the shorter mRNAs used in this study compared with that of other species. However, the average or median transcript length of locust lncRNAs is also longer than that of these species. The reason for the longer transcript length of locust lncRNAs compared with that of other species needs to be further studied. The large portion of noncoding region in the locust possibly contributes to this phenomenon to a certain extent [24]. On the other hand, locust lncRNAs differ significantly from mRNAs in terms of structure and expression. Locust lncRNAs possess longer exons, fewer exons per transcript, lower global expression level than mRNAs, and more lincRNAs, which is similar to other species, such as flies [5,34], zebrafish [3], and humans [1]. These results indicate the particular and common structure characteristics of locust lncRNAs with other species. The characteristics of longer exon may influence the evolution, alternative splicing and expression of locust lncRNAs.

### Interactions of lncRNA and mRNAs in response to phase change of locusts

The interaction analysis of lncRNAs and mRNAs in time-course treatments demonstrated that more lncRNAs were expressed in gregarious locust than in solitarious ones, and higher percentage of lncRNAs were specifically expressed in the two locust phases than mRNAs. One interesting phenomenon is that lncRNAs respond more rapidly to crowding and isolation than mRNAs. The quicker responses were reflected by the higher ratio of lncRNAs regulated at the 4 h time point and their faster expression change rate in the first 4 h. Therefore, both the ratio and the expression change rate of early changed lncRNAs are notably higher than those of early changed mRNAs, indicating that locust lncRNAs are more sensitive to population density changes than mRNAs. The initial role of lncRNAs in the transcriptional process is extremely crucial in the response to the changes in population density, displaying the similarity with the initial role of *CSP* gene in the locust phase change [25]. Given that lncRNAs can regulate transcription, translation, and stability of protein-coding genes [7], it is reasonable for lncRNAs responding firstly as regulators. This finding may be explained by the low stability and rapid turnover rate of lncRNAs [35]. However, the transcription speed of lncRNAs may be more important than the other reasons above [36]. The density-cued expression of LNC1010057 could be regulated by several transcription factors such as ZNF300, lolal and Gtf2e2 which were co-expressed with LNC1010057 (Table S6). The lncRNA expressions in the time series of the development and other biological processes have been fully studied in other species [5,37]; however, the dynamic expression of lncRNAs in time-course treatments on behaviors has been rarely described in insects. The temporal analysis uncovered for the first time the initial roles of lncRNAs in response to the change in population density. Thus, the rapid response of noncoding RNAs may help organisms rapidly adapt to the changes in environmental conditions.

### Identification and functional verification of hub lncRNAs

Given that functions of lncRNAs can be inferred by protein-coding genes in co-expression networks and nearby genes in the genome [9], we performed co-expression analysis to characterize the potential functions of early changed lncRNAs instead of the nearby genes. Because the nearby genes which changed expressions were extremely few to analyze (Table S7). Several lncRNAs are strongly associated with known phase-related genes, such as *NPYR* and *NPF1a* genes in the NO_2_ pathway [27,28], and *Ebony* and *Vat1* genes on the dopamine pathway [26], which have been confirmed closely linked with the locust phase change. Thus, this result provides a new noncoding layer that is connected to the primary mechanism, in which protein-coding genes participated in the locust phase change. At the same time, functional annotation indicates that the locust phase change is involved in multiple regulatory mechanisms and concentrates on the common pathways, as reported by our previous studies [25,26,28].

In the present study, we predicted two hub and homologous lncRNAs as potential regulators because they were conversely expressed in crowding and isolation. LNC1010057 but not LNC992414 was validated to primarily participate in the locust phase change by experiment in vivo, although they shared several common domains. The LNC992414 knockdown was insufficient to cause the shift from the gregarious phase to the solitarious phase. Thus, LNC1010057 is a repetitive element-containing lncRNA which regulates the locust behavior, although it is not conserved among other species. In other species, repetitive element-containing lncRNAs have also been proved to have functions. In primates, an Alu element-inserted lncRNA 5S-OT regulates the alternative splicing of target genes through complementary base pairs [38]. Another SINE-containing lncRNA regulates myogenesis in rodents [39]. In the nervous system, the lncRNA and repetitive element expressions were bioinformatically predicted to function in synaptic plasticity, but no experimental evidence proved this proposal [40]. In migratory locust, the repetitive elements constitute more than the half of the genome. The present study offers important evidences about their important functions in the phase-related behavior change in locusts. On the other hand, various roles of homologous lncRNAs may be due to the extremely high relative expression level of LNC1010057 offsetting the decrease in LNC992414 expression level. Another explanation for this result is that the secondary structure of the longer lncRNA LNC992414 blocks the binding sites for targets to function with [41,42]. The diverse functions of homologous lncRNAs are also identified for lncRNA SNHG10 and SCARNA13 in hepatocarcinogenesis [43]. Finally, given that no neighboring genes within 100 Kb were found for LNC1010057 in the locust genome, LNC1010057 showed the least possibility for *cis-*regulation. Therefore, LNC1010057 may regulate distal target genes through *trans*-acting mechanism [44]. The genes co-expressed with LNC1010057 in CS and IG networks may be the *trans*-targets that should be validated in the future. However, we cannot exclude the possibility that the neighboring genes which are more than 100kb faraway may interact with LNC1010057. With the development of sequencing quality, there may be neighboring genes that can be candidates for further study as well.

In conclusion, this work discovered the relationships between protein-coding genes and lncRNAs in the locust phase change, proved the regulatory role of lncRNAs in phase-related behavior in insects, and provided additional insights into the mechanism underlying the phenotype plasticity of animals.

## Materials and methods

### Insects

Gregarious and solitary locusts used in experience were obtained from the same locust colony and maintained at the laboratory. Gregarious locusts were cultured in well-ventilated cages (40 cm × 40 cm × 40 cm) at a density of 500–1,000 locusts per cage for eight generations. Solitary locusts were cultured individually in white metal boxes (10 cm × 10 cm × 25 cm) supplied with fresh air for more than 10 generations. Both colonies were maintained under the same photocycle regime and temperature conditions and fed with the same food as described in the previous works [29].

### Isolation and crowding treatments

For isolation treatment, the gregarious locusts were separately raised as solitary locusts as described above. After 4, 8, or 16 h of isolation, the brains of the third-day fourth-instar gregarious locusts were collected at 2 p.m. of the day. The gregarious locust brains were collected as the 0 h time point sample for the isolation course. For crowding treatment, 10 fourth-instar solitarious locusts were reared together with a stimulating group of 20 fourth-instar gregarious locusts in a small cage (10 cm × 10 cm × 10 cm). A total of 6 replicate cages were prepared for each time point of crowding treatment. After 4, 8, or 16 h of crowding, the brains of the third-day fourth-instar solitarious locusts were collected at 2 p.m. of the day. The solitarious locust brains were collected as the 0 h time point sample for the crowding course. All locust samples from isolation and crowding treatments were immediately frozen using liquid nitrogen for storage. Three biological replicates samples were collected at the same time point. Each replicate contained 20 insects, with equal numbers of males and females.

### RNA purification, cDNA library construction, and RNA-seq

The total RNA was extracted from the collected locust brains by the TRIzol reagent (Cat. No. 15596-026, Invitrogen, Carlsbad, CA) according to the protocol. The quantity and purity of the total RNA extracted as described above were determined with the RIN value of >7 by ND-1000 Nanodrop (Thermo Fisher Scientific, Wilmington, DE) and Agilent 2200 Tapestation (Agilent, Waldbronn, Germany). RNA integrity was examined by agarose gel electrophoresis. Then, ribosomal RNA was removed from total RNA using EpicentreRibo-Zero rRNA Removal Kit, (Cat. No. MRZH11124, Illumina company, San Diego, CA). rRNA-depleted RNA was randomly fragmented into short segments and instantly used as templates to synthesize the first cDNA by random hexamers. Next, the second cDNA was synthesized, and RNA was digested by RNase H. Then the double-stranded DNA was purified and added with bases A and adaptor in the 3′-end. After selection by AMPureXP beads, the second strand cDNA containing U was degraded by the enzyme USEB. Finally, the cDNA library for sequencing was constructed by PCR amplification. Then, the quality of the cDNA library was also inspected, and sequencing was performed on an Illumina Hiseq3000/2500. In the end, the sequencing reads was produced with the length of pair-ended to be 2×150 bp (PE150).

### Transcriptome assembly and lncRNA identification

Prior to assembly, the quality of raw reads was evaluated by FastQC and cleaned by removing adaptor sequences and poor-quality reads. Then, the clean data were mapped to the *Locusta migratoria* genome (version 2.4.1, available at http://159.226.67.243/) using the read aligner HISAT2 (version 2.1.0). Next, the transcriptome was assembled by the StringTie (version 1.3.1) on the basis of the reads mapped to the reference locust genome. The assembled transcripts were annotated by the Cuffcompare program from the Cufflinks package. lncRNA screening and an analysis pipeline was developed as follows. (1) The known protein-coding transcripts according to the annotation of the reference genome were discarded. (2) The transcripts with the length of < 200 bp, single exon, and FPKM of < 0.1 were filtered out. (3) The transcripts with class code “=“, “j”, “p”, “s” were excluded. (4) The transcripts with protein coding potential (Coding-Potential Assessment Tool (CPAT) score > 0.38, Coding Non-coding Index (CNCI) score > 0 and Coding Potential Calculator (CPC) score > 0) were removed [45–47]. (5) The transcripts with similarity to known protein-coding domains in the Pfam (version 31.0) database (E-value < 1e-3) were eliminated. (6) The transcripts with FPKM of < 1.0 were discarded. Finally, the remaining transcripts were classified as lncRNAs.

### Differential expression analysis of lncRNAs and mRNA

The FPKMs of all transcripts in each sample were calculated by StringTie (version 1.3.1). The differential expression analysis of lncRNA and mRNAs between two conditions was performed using the DESeq2 (version 1.10.1) package in the R project. The transcripts with *P* < 0.05 and absolute fold change > 2.0 were considered differentially expressed.

### qRT-PCR

qRT-PCR was performed to validate the expression of candidate lncRNAs and target genes selected from the bioinformation prediction. For each sample, 2 µg total RNA was reverse-transcribed to cDNA with random primers by Moloney murine leukemia virus reverse transcriptase (Cat. No. M1705, Promega, Madison, WI). Then, quantitative PCR was performed using the SYBR Green I Master (Cat. No. 4887352001, Roche, Mannheim, Germany) on a Light Cycler 480 instrument (Roche, Mannheim, Germany) to evaluate the expression level. Six independent biological replicates were used for each condition in expression analysis. Then, the relative expression levels of transcripts were calculated through the 2^-ΔΔCt^ method with RP49 as the endogenous reference gene. The melting curve was used to validate unique amplification, and all the qPCR amplifications were sequenced to confirm the specific products. The absolute expression levels of lncRNAs were calculated in accordance with the standard curves which were derived from the absolute quantitative PCR. The standard curves presented the correlation between the Ct values and real RNA concentrations. Table S8 lists the primers used.

### Temporal specificity analysis

We used index τ to measure the specific expression [48] as follows:

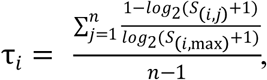

where *n* is the number of samples tested, *S*_*(i,j)*_ is the FPKM of transcript *i* in sample *j*, and *S*_*(i,max)*_ is the highest FPKM of the transcript *i* in *n* samples.

### Time series profile analysis of transcript expression

The DE transcripts in crowding and isolation were used to perform the expression pattern analysis by STEM (version 1.3.11). We selected a set of 26 distinct temporal expression profiles independent of the data to summarize the transcript expression change independent of the data. The expressions of transcripts were normalized (log2(0h/0h), log2(4h/0h), log2(8h/0h), log2(16h/0h)) before being clustered. lncRNAs or mRNAs were assigned to the model profiles that most closely represented their expression patterns, as determined by the correlation coefficient (*r* > 0.7). The significance levels at which the number of transcripts assigned to a model profile compared to the expected number of transcripts assigned were calculated for each model profile. The profiles with *P* < 0.05 were identified as significant temporal expression profiles which obviously responded to the treatment.

### LncRNA–mRNA Co-expression network construction

The correlation coefficients of lncRNAs and mRNAs were evaluated based on the normalized expression (FPKM), as determined by using the Pearson correlation method in R software. The expression levels of the transcripts at each time point was computed by averaging the FPKMs in three biological replications. The protein-coding genes with correlation coefficients of > 0.9 (positive) or < -0.9 (negative) with the corresponding lncRNAs were considered co-expressed genes. These correlations between lncRNAs and mRNAs were put into the Cytoscape software (v3.6.1) to constructed co-expression network.

### Functional pathway analysis

KEGG enrichment analyses were performed using the Enrichment widget implemented in the Locust Mine database (http://www.locustmine.org:8080/locustmine) [49]. Pathway and biological process with *P* < 0.05 were regarded as enriched.

### Rapid amplification of cDNA ends (RACE)

The 5′ and 3′ RACE experiments were performed following the instruction manual (Cat. No. 634923, SMARTer RACE cDNA Amplification Kit, Clontech, Mountain View, CA). The nest RACE was performed on the templates that were diluted 50 times. The touchdown PCR programs were used to amplify specific target sequences. Table S8 shows the primers used.

### RNAi

First, the dsRNA of the *GFP* and lncRNA for RNAi were prepared using T7 RiboMAX Express RNAi system (Cat. No. P1700, Promega, Madison, WI). Then, 69 nl dsRNA for each treatment was injected into the middle of the brain of each locust through microinjection. After injection, the gregarious locusts were returned to the cage for rearing. Every treatment included six biological repeats with 6–7 locusts. After 3 days, the brains of the locusts were sampled, followed by RNA purification and qRT-PCR to confirm the effectivity of RNAi.

### Behavior test in arena

Behavioral assay was conducted in a rectangular perspex arena (40 cm × 30 cm × 10 cm) with opaque walls in accordance with a previous study [25]. Two similar champers (7.5 cm × 30.0 cm × 10.0 cm) were on each side of the arena, with one containing 30 fourth-instar gregarious nymphs and the other being empty. The locusts were gently placed into the assay arena through a cylinder beneath it and recorded for 300 s after visual appearance. The video recording and behavioral data analysis were performed using an EthoVision video tracking system (Noldus, Wageningen, The Netherlands).

Five different behavioral parameters, including attraction index (AI), duration in the attraction zone, duration in the repulsion zone, TDMV, and TDM were extracted and regarded as the categorical behavior markers. The behavioral phenotypes of locusts were analyzed by applying the binary logistic regression model that was described in our previous study [25]. The regression model was described as follows: P_*greg*_ = e^*η*^/ (1 + e^*η*^); *η* = -2.11 + 0.005 × AI + 0.012 × TDM + 0.015 × TDMV, where P_*greg*_ represents the probability of a locust being regarded as the gregarious phase (P_*greg*_ = 1 indicates that the locust is fully gregarious, whereas P_*greg*_ = 0 indicates fully solitarious behavior), AI = total duration in attraction zone (near to camper containing locusts) - total duration in repulsion zone (near to empty camper); this parameter indicated the extent to which the tested locusts were attracted by the gregarious locusts, and TDMV and TDM indicate the locomotor activity levels. In the behavioral assay, more than 29 locust individuals were tested for each treatment.

### Statistical Analysis

The ks-test was conducted to analyze the differences between lncRNA and mRNA in length and exon/intron number. Pearson’s Chi-squared test was performed to compare the percentages of differentially changed lncRNAs and mRNAs. Student’s *t*-test was used to analyze the effectiveness of RNAi and the changes in specific parameters in arena assay. The data analysis of behavioral phase change in the arena assay was performed by Mann–Whitney U test for their non-normal distribution characteristics.

### Data availability

The Fastq files of the strand-specific transcriptome sequence for 24 samples in time course have been deposited in the Genome Sequence Archive [50] in BIG Data Center, Beijing Institute of Genomics (BIG), Chinese Academy of Sciences, under accession numbers CRA002379, CRA002379 that are publicly accessible at https://bigd.big.ac.cn/bioproject/browse/PRJCA002298.

## Author Contributions

LK supervised the project and conceived the study. TL performed most of the bioinformatics analysis and experiments, and wrote the paper. BC took part in the discussion and comments in the manuscript draft. PY contributed the analytic tools. DW and BD helped with data analyses. All authors read and approved of the final manuscript.

## Acknowledgements

We thank Dr. Runsheng Chen, Dr. Fangqing Zhao, Dr. Shunmin He, and Dr. Dongdong Zhang for their helpful comments. We thank Dr. Feng Jiang, Dr. Jing He and Dr. Ding Ding for their valuable discussion. We thank Dr. Xia Zhang and Jing Wang for technical help. We are grateful to Ya′nan Zhu for locust rearing. We thank ShineWrite company for the help in language polishing. This study was supported by the National Science Foundation of China (grant no. 31430023, 31872304 and 31771452).

## Competing interest

The authors declare no competing financial interests.

## Supplementary material

### Supplementary Figures

**Figure S1 The workflow of lncRNA identification**

**A**. Pipeline of lncRNA identification. The numbers on the right represent the obtained transcript numbers in the process. **B**. Saturation test of sequencing data. **C**. Venn diagram of the number of lncRNA predicted by using CNCI, CPC, CPAT, and by searching the pfam database. The numbers in brackets represent lncRNAs with FPKM > 1 and other numbers represent lncRNAs with FPKM > 0.1.

**Figure S2 Correlation of the lncRNA expression between qPCR and RNAseq**

The x- and y-axis represent the relative expression of lncRNAs (log(v(i)/v(0))) evaluated in RNAseq and qPCR, respectively. V(i) and v(0) represent the expressions of lncRNA at i h (i = 4, 8, 16) time point and 0 h time point, respectively.

**Figure S3 Different expressions of lncRNAs and mRNAs in CS and IG**

**A**. The number of totally and specifically expressed lncRNAs and mRNAs at each time point of time course in CS and IG. **B**. The total number of DE lncRNAs and mRNAs in CS and IG. **C**. The number of DE lncRNAs and mRNAs at CS16h relative to G, and at IG16h relative to S.

**Figure S4 Hierarchical clustering of DE lncRNAs and mRNAs**

The hierarchical clustering of DE lncRNAs and mRNAs in CS (**A**) and IG (**B**). The color bars on the right represent the correlations between samples. The correlations were calculated by the Pearson correlation test.

**Figure S5 Expression patterns of DE mRNAs and lncRNAs in CS and IG were analyzed via STEM**

Expression patterns clustered for lncRNAs and mRNAs in CS (**A**) and IG (**B**). A total of 26 profiles independent of expression data were set to summarize the expression change patterns. The numbers in the left upper part of boxes are profile serial numbers, those in the left lower part are p-values, and those in the right lower part are numbers of transcripts contained in profiles. The colored boxes are the profiles that showed significant p-values (*P* < 0.05). The significant profiles were colored randomly by the software.

**Figure S6 Similarity analysis of repetitive elements of LNC1010057 and LNC992414**

**A.** Sequence alignment of repetitive element A of LNC1010057 and LNC992414. Identical residues are colored in red (A residue), orange (T residue), green (G residue) and blue (C residue). Dashes represent gaps in the sequence relative to counterparts in the alignments. **B**. Sequence alignment of repetitive element B of LNC1010057 and LNC992414. **C**. Sequence alignment of repetitive element C of LNC1010057 and LNC992414.

**Figure S7 Changes in other phase-related behavior parameters after RNAi of repetitive elements and LNC992414**

**A**. Change in movement speed of gregarious locusts after repetitive elements knockdown. **B**. Changes in duration of gregarious locusts in the attraction and repulsive zone after repetitive elements knockdown. **C**. Changes in movement duration of gregarious locusts in border and central zone after repetitive elements knockdown. **D**. Changes in turn times and mean turn angle of gregarious locusts after repetitive elements knockdown. **E**. Change in movement speed of gregarious locusts after LNC992414 knockdown. **F**. Changes in duration of gregarious locusts in the attraction and repulsive zone after LNC992414 knockdown. **G**. Changes in movement duration of gregarious locusts in border and central zone after LNC992414 knockdown. **H**. Changes in turn times and mean turn angle of gregarious locusts after LNC992414 knockdown. Measurements are shown as mean ± SE. \**P <* 0.05, \*\**P <* 0.01, \*\*\**P <* 0.001 (Student’s *t*-test).

### Supplementary Tables

**Table S1 Clean data quality (Q30) for each sample**

**Table S2 Summary of lncRNA and protein-coding RNAs in total transcripts**

**Table S3 *In-cis* target genes of DE lncRNAs in S vs G locusts**

**Table S4 Pearson correlation coefficient between early changed lncRNAs and all mRNAs in CS and IG**

**Table S5 Correlations between known phase-related genes and lncRNAs**

**Table S6 Transcription factors co-expressed with LNC1010057**

**Table S7 The neighboring genes of early changed lncRNAs**

**Table S8 Primers used in this study**

## Notes

### Competing Interest Statement

The authors have declared no competing interest.

