## Supplementary text for "Long Noncoding RNA Derived from LncRNA–mRNA Co-expression Networks Modulates the Locust Phase Change"

^*^ Corresponding author.

**Running title**: *Li T / LncRNA Regulates the Locust Phase Change*

^a^ORCID: 0000-0002-5503-5396.

^b^ORCID: 0000-0002-1238-3948.

^c^ORCID: 0000-0001-5496-8357.

^d^ORCID: 0000-0003-2764-5477.

^e^ORCID: 0000-0003-4197-818X.

^f^ORCID: 0000-0003-4262-2329.

**Supplementary Figures:**

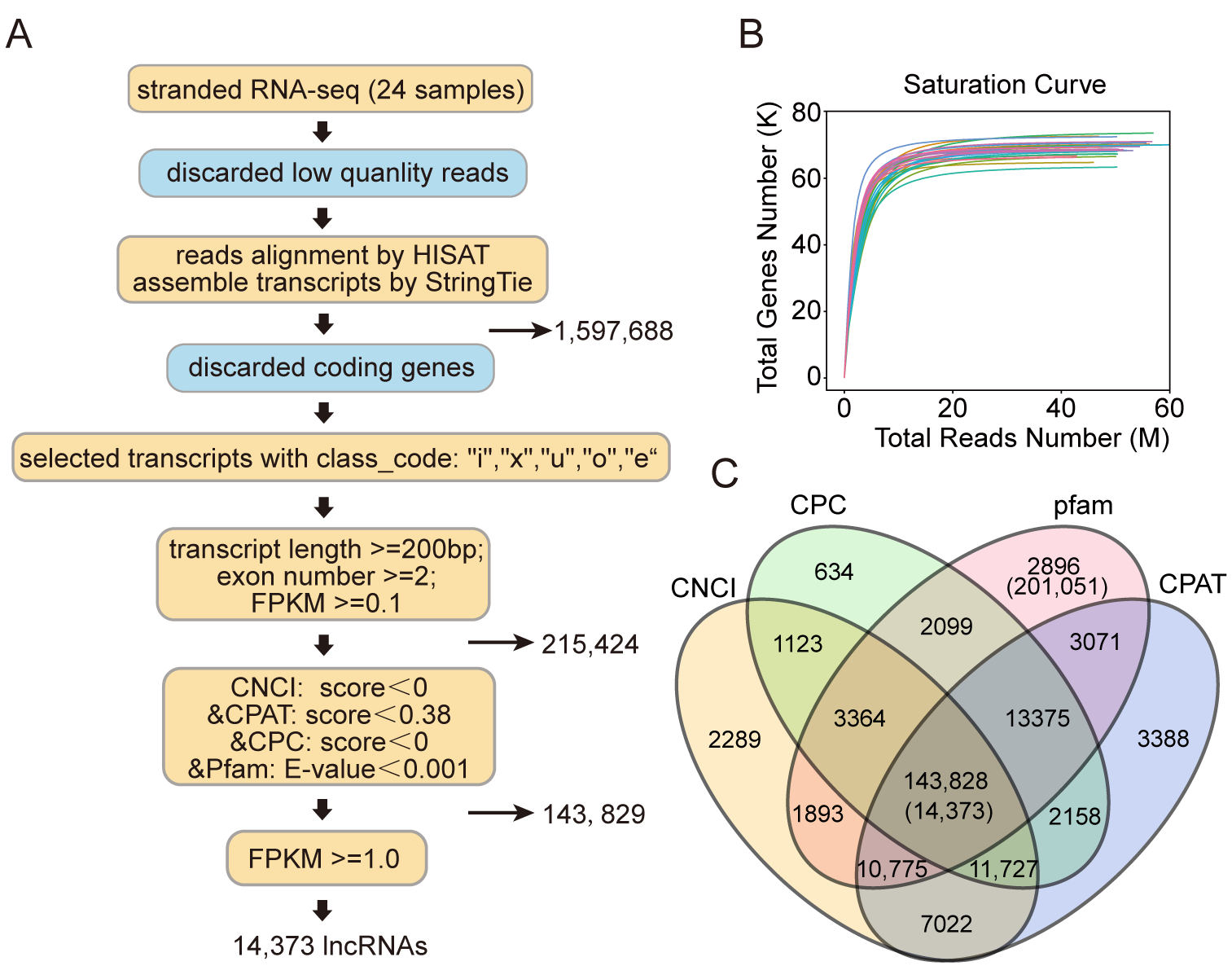

**Figure S1 The workflow of lncRNA identification**

**A**. Pipeline of lncRNA identification. The numbers on the right represent the obtained transcript numbers in the process. **B**. Saturation test of sequencing data. **C**. Venn diagram of the number of lncRNA predicted by using CNCI, CPC, CPAT, and by searching the pfam database. The numbers in brackets represent lncRNAs with FPKM > 1 and other numbers represent lncRNAs with FPKM > 0.1.

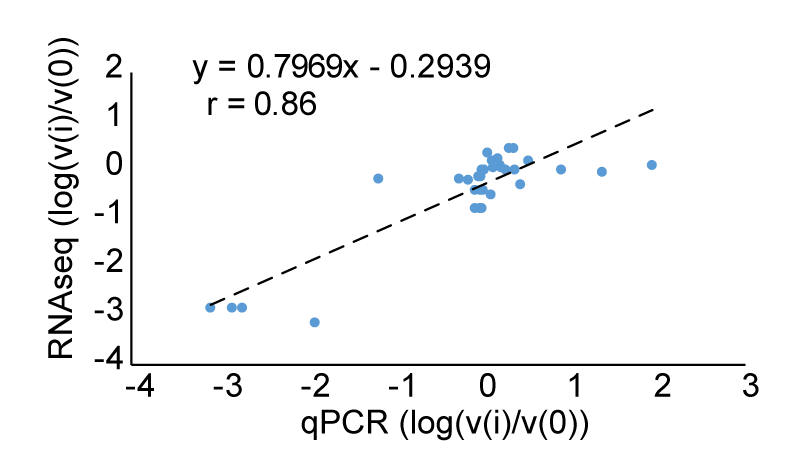

**Figure S2 Correlation of the lncRNA expression between qPCR and RNAseq**

The x- and y-axis represent the relative expression of lncRNAs (log(v(*i*)/v(0))) evaluated in RNAseq and qPCR, respectively. V(*i*) and v(0) represent the expressions of lncRNA at *i* h (*i* = 4, 8, 16) time point and 0 h time point, respectively.

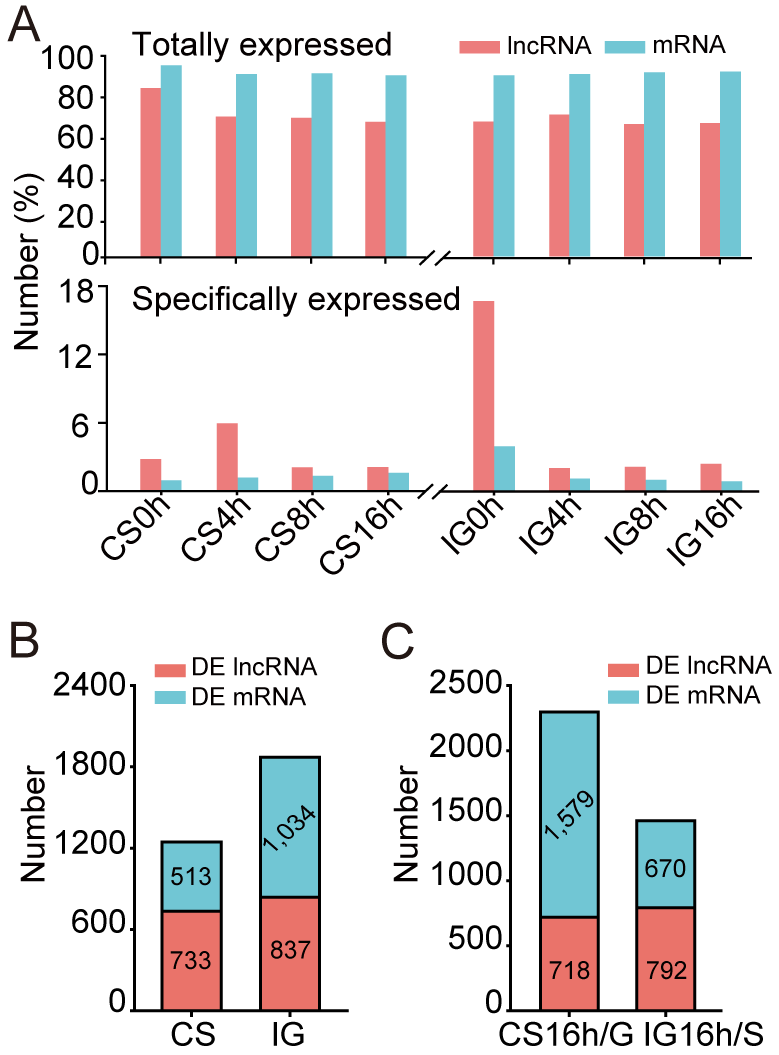

**Figure S3 Different expressions of lncRNAs and mRNAs in CS and IG**

**A**. The number of totally and specifically expressed lncRNAs and mRNAs at each time point of time course in CS and IG. **B**. The total number of DE lncRNAs and mRNAs in CS and IG. **C**. The number of DE lncRNAs and mRNAs at CS16h relative to G, and at IG16h relative to S.

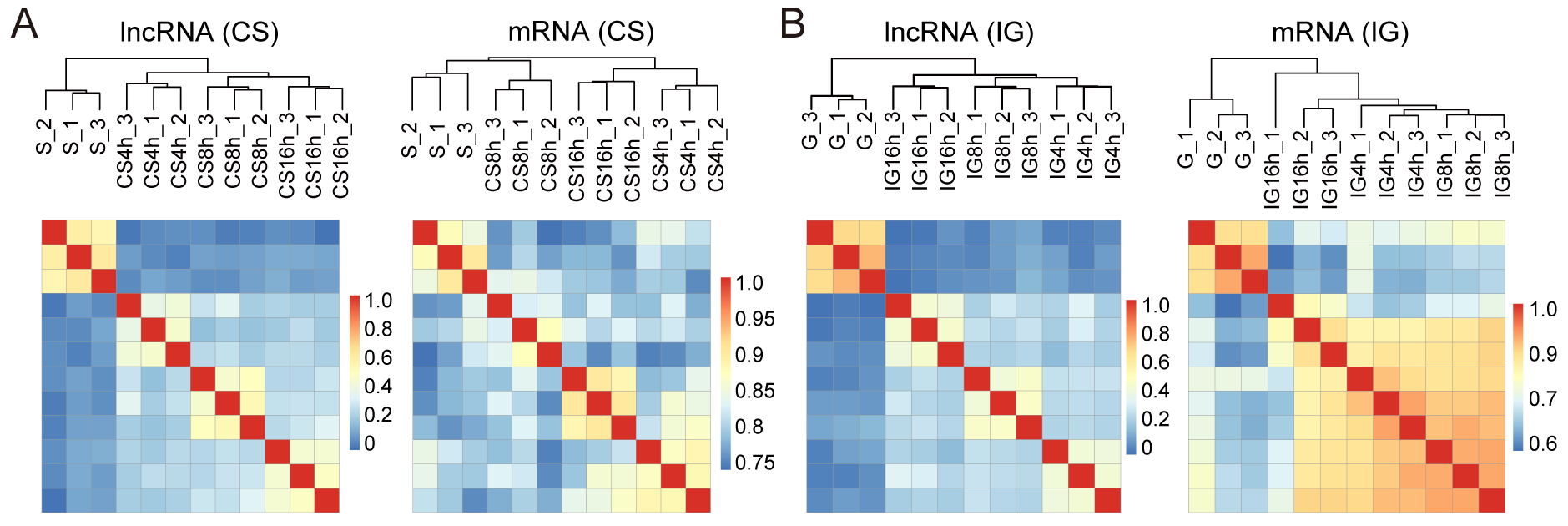

**Figure S4 Hierarchical clustering of DE lncRNAs and mRNAs**

The hierarchical clustering of DE lncRNAs and mRNAs in CS (**A**) and IG (**B**). The color bars on the right represent the correlations between samples. The correlations were calculated by the Pearson correlation test.

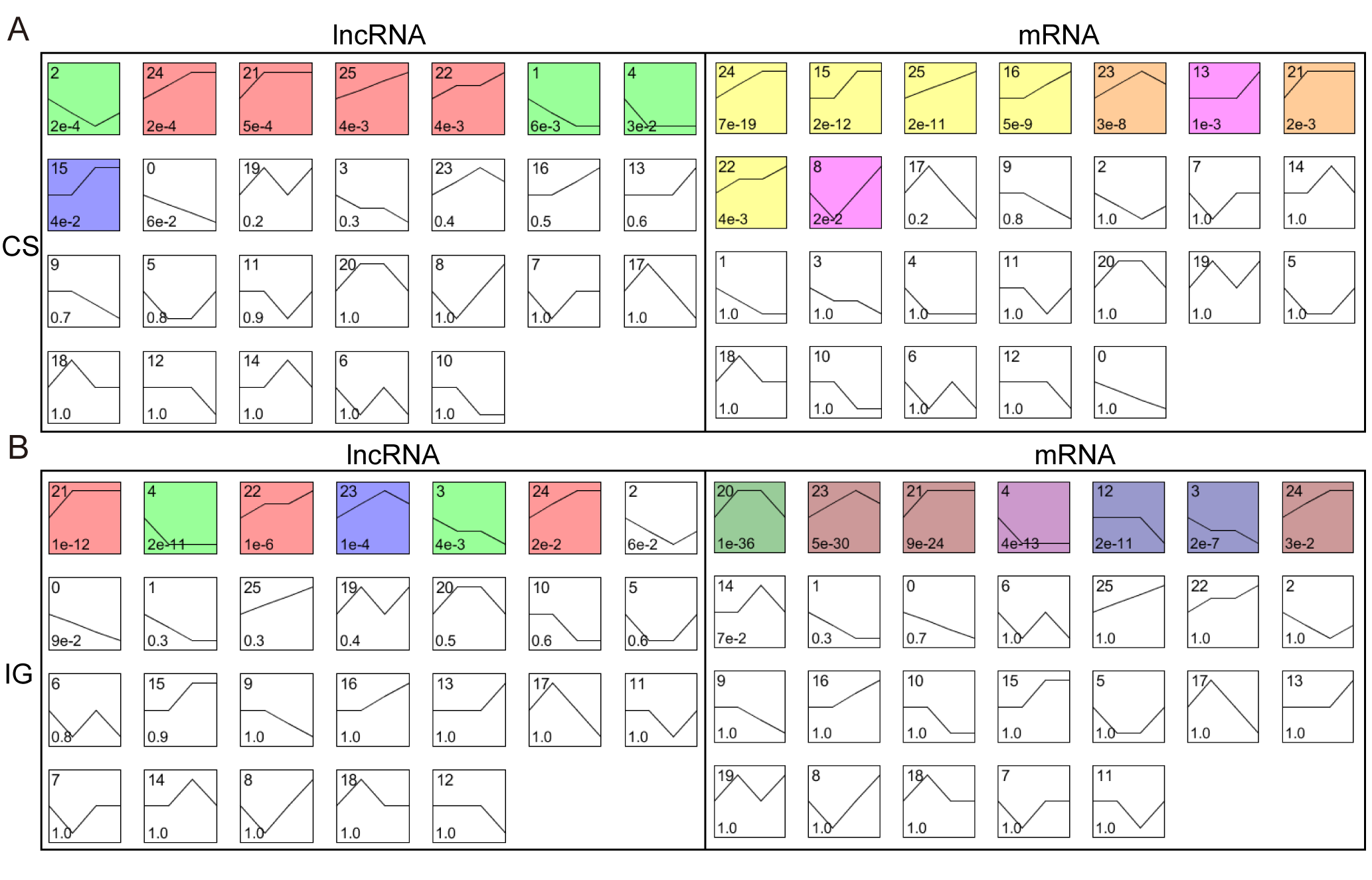

**Figure S5**  **Expression patterns of DE mRNAs and lncRNAs in CS and IG were analyzed via STEM**

Expression patterns clustered for lncRNAs and mRNAs in CS (**A**) and IG (**B**). A total of 26 profiles independent of expression data were set to summarize the expression change patterns. The numbers in the left upper part of boxes are profile serial numbers, those in the left lower part are p-values, and those in the right lower part are numbers of transcripts contained in profiles. The colored boxes are the profiles that showed significant p-values (*P* < 0.05). The significant profiles were colored randomly by the software.

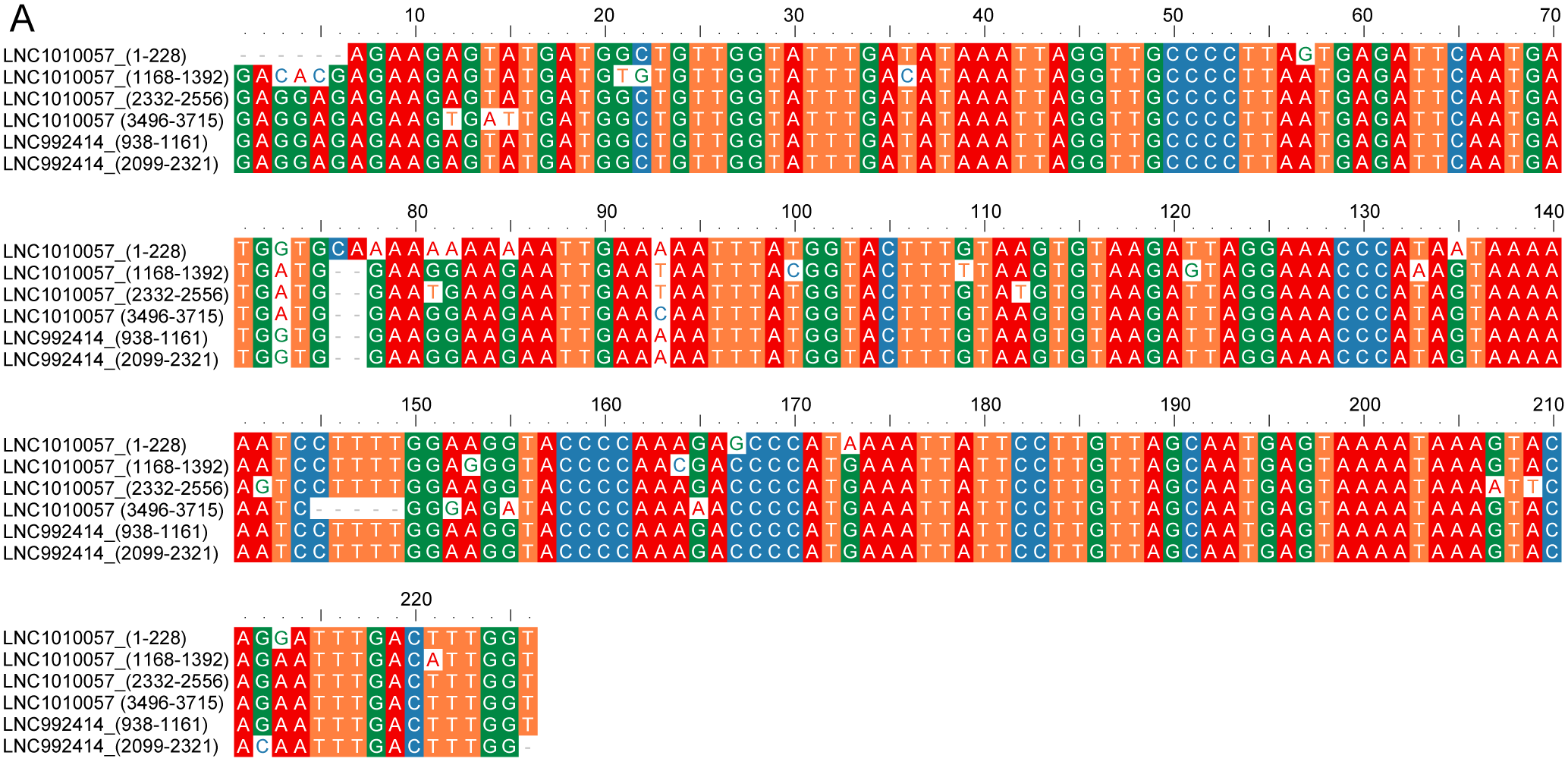

**Figure S6 Similarity analysis of repetitive elements of LNC1010057 and LNC992414**

**A.** Sequence alignment of repetitive element A of LNC1010057 and LNC992414. Identical residues are colored in red (A residue), orange (T residue), green (G residue) and blue (C residue). Dashes represent gaps in the sequence relative to counterparts in the alignments.

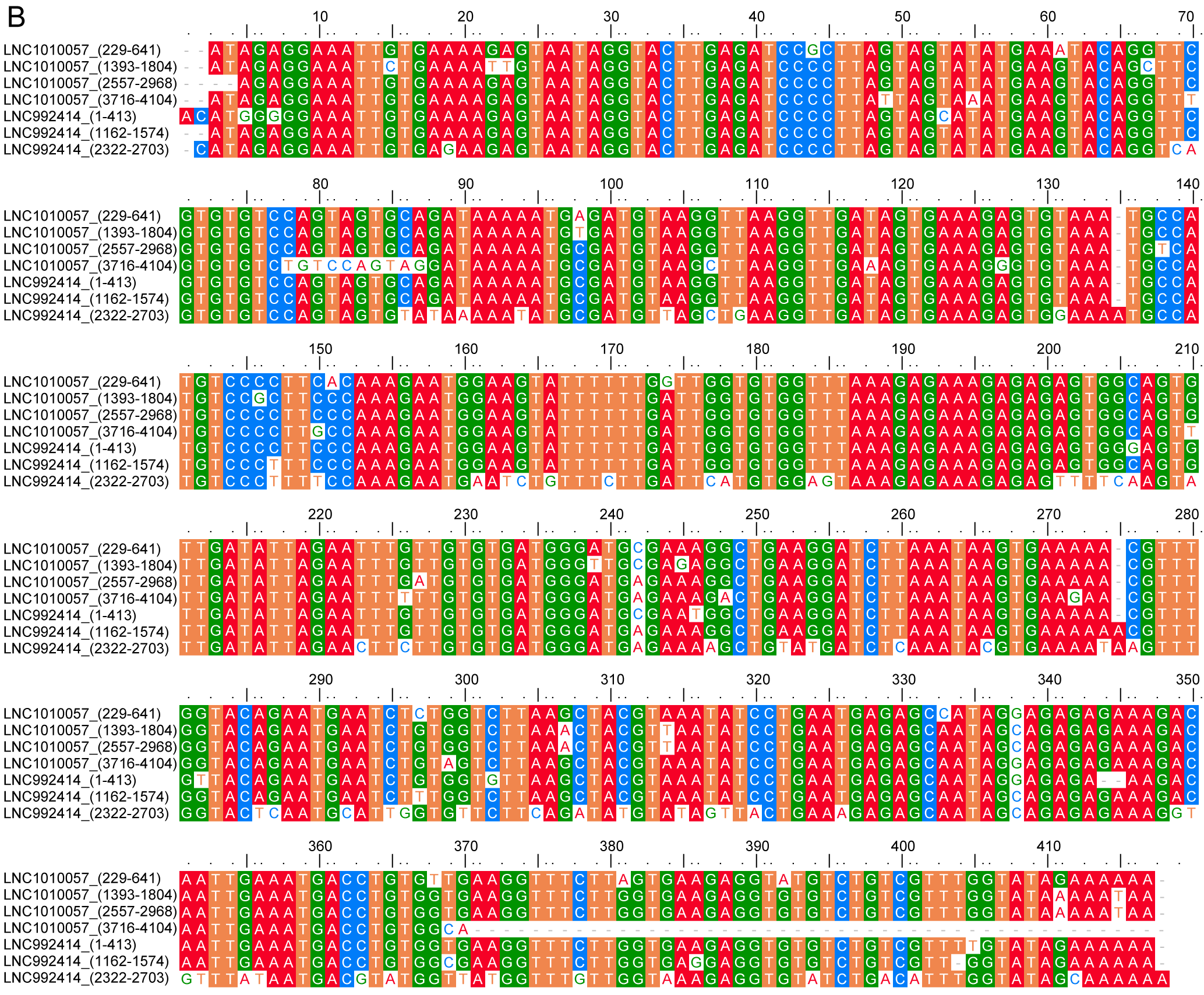

**Figure S6 Similarity analysis of repetitive elements of LNC1010057 and LNC992414**

**B**. Sequence alignment of repetitive element B of LNC1010057 and LNC992414.

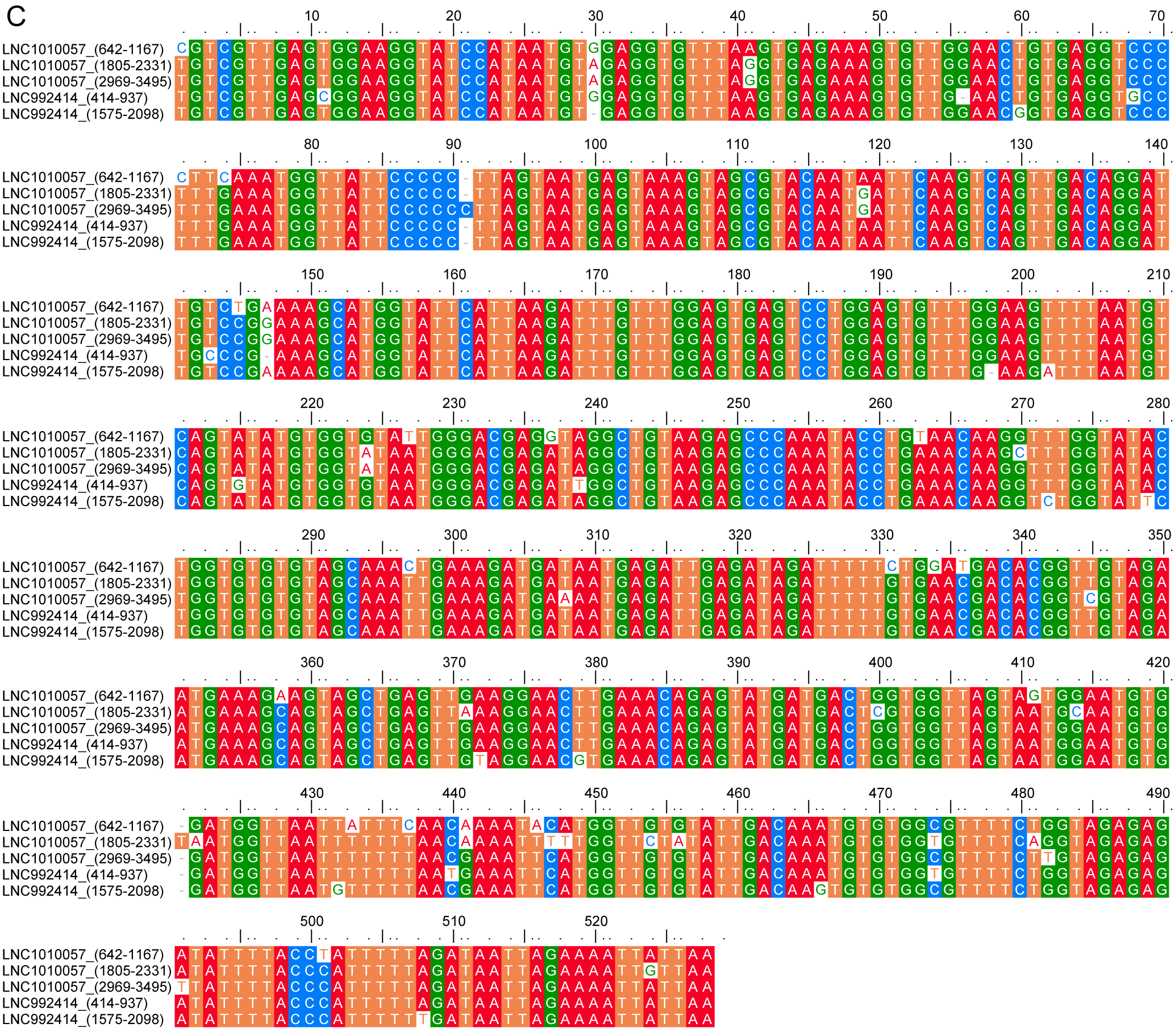

**Figure S6 Similarity analysis of repetitive elements of LNC1010057 and LNC992414**

**C**. Sequence alignment of repetitive element C of LNC1010057 and LNC992414.

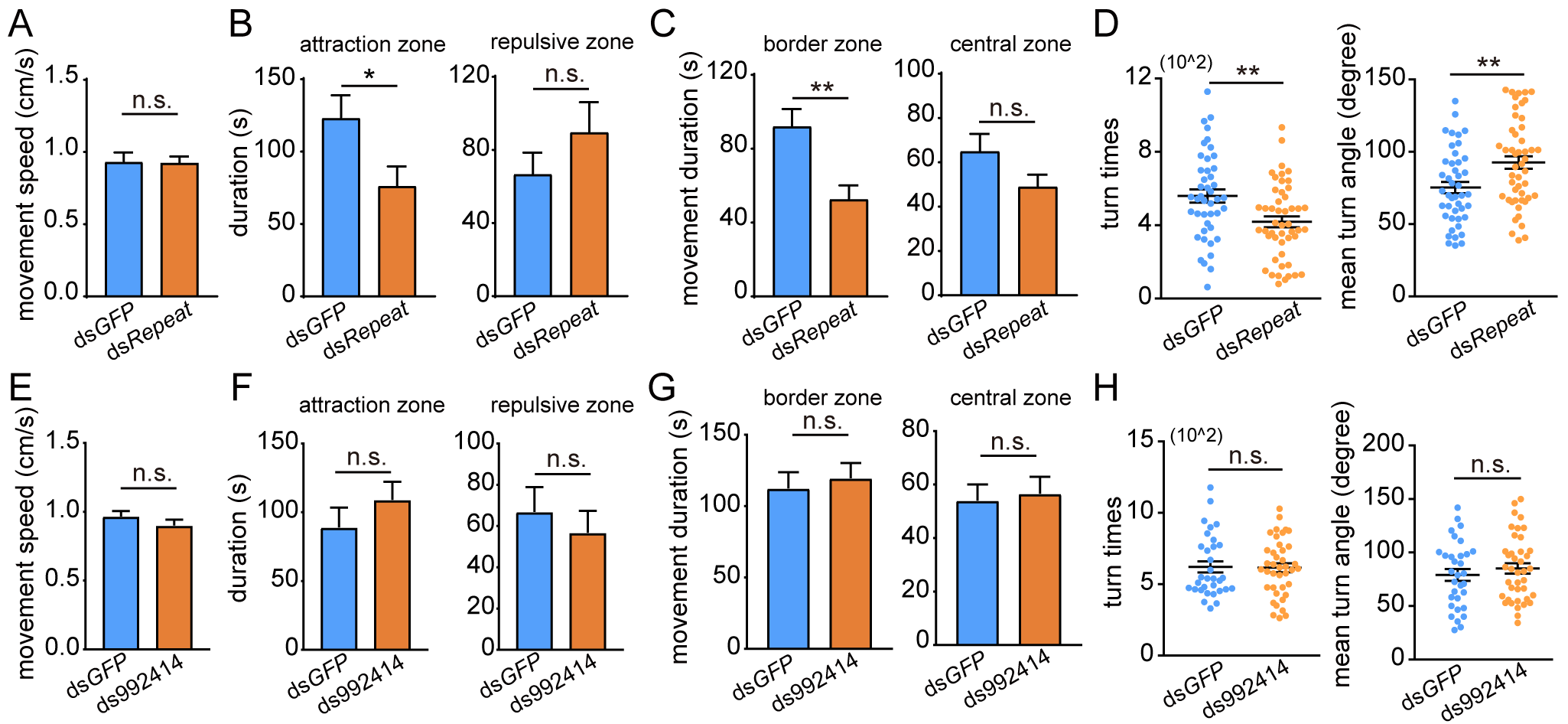

**Figure S7 Changes in other phase-related behavior parameters after RNAi of repetitive elements and LNC992414**

**A**. Change in movement speed of gregarious locusts after repetitive elements knockdown. **B**. Changes in duration of gregarious locusts in the attraction and repulsive zone after repetitive elements knockdown. **C**. Changes in movement duration of gregarious locusts in border and central zone after repetitive elements knockdown. **D**. Changes in turn times and mean turn angle of gregarious locusts after repetitive elements knockdown. **E**. Change in movement speed of gregarious locusts after LNC992414 knockdown. **F**. Changes in duration of gregarious locusts in the attraction and repulsive zone after LNC992414 knockdown. **G**. Changes in movement duration of gregarious locusts in border and central zone after LNC992414 knockdown. **H**. Changes in turn times and mean turn angle of gregarious locusts after LNC992414 knockdown. Measurements are shown as mean ± SE. **P <* 0.05, ***P <* 0.01, ****P <* 0.001 (Student’s *t*-test).

**Supplementary Tables:**

**Table S1** **Clean data quality (Q30) for each sample**

| **Sample ID** | **Read Sum** | **Base Sum** | **GC(%)** | **Q30(%)** |
| --- | --- | --- | --- | --- |
| S-1 | 65,202,185 | 19,201,909,451 | 44.86 | 94.53 |
| S-2 | 46,022,828 | 13,469,126,574 | 47.26 | 94.83 |
| S-3 | 49,433,606 | 14,564,010,885 | 44.37 | 95.04 |
| CS4H-1 | 56,993,730 | 16,767,978,060 | 48.24 | 94.84 |
| CS4H-2 | 50,427,966 | 14,816,869,841 | 44.74 | 94.85 |
| CS4H-3 | 50,318,526 | 15,095,557,800 | 45.29 | 91.1 |
| CS8H-1 | 50,363,248 | 14,924,691,342 | 44.0 | 94.71 |
| CS8H-2 | 60,118,222 | 17,786,205,431 | 44.77 | 94.74 |
| CS8H-3 | 52,104,206 | 15,337,621,145 | 44.97 | 95.17 |
| CS16H-1 | 50,141,494 | 14,825,288,188 | 46.11 | 94.94 |
| CS16H-2 | 48,694,324 | 14,402,782,691 | 43.78 | 95.08 |
| CS16H-3 | 50,142,156 | 14,831,798,260 | 44.7 | 94.8 |
| G-1 | 42,891,678 | 12,466,904,814 | 44.37 | 92.87 |
| G-2 | 56,304,618 | 16,314,356,487 | 42.5 | 94.95 |
| G-3 | 46,949,291 | 13,589,981,414 | 42.05 | 92.67 |
| IG4H-1 | 54,497,032 | 15,780,883,888 | 42.97 | 95.25 |
| IG4H-2 | 50,615,150 | 14,722,016,243 | 42.97 | 95.08 |
| IG4H-3 | 56,799,204 | 16,574,197,519 | 42.45 | 94.95 |
| IG8H-1 | 51,435,271 | 14,938,545,678 | 42.76 | 95.15 |
| IG8H-2 | 47,121,268 | 13,708,374,232 | 42.49 | 95.17 |
| IG8H-3 | 42,813,933 | 12,481,691,994 | 42.27 | 95.4 |
| IG16H-1 | 55,691,716 | 16,252,214,016 | 41.93 | 95.5 |
| IG16H-2 | 50,275,776 | 14,715,933,534 | 41.4 | 94.76 |
| IG16H-3 | 53,237,615 | 15,570,056,972 | 42.16 | 95.52 |

**Table S2** **Summary of lncRNA and protein-coding RNAs in total transcripts**

| **Transcripts** | **LncRNA** | **Coding** |
| --- | --- | --- |
| No. of sequence | 14,373 | 15,309 |
| No. of bases | 38,081,213 | 20,259,011 |
| Max sequence length (bp) | 25,251 | 22,501 |
| Average sequence length (bp) | 2649 | 1152 |
| Media sequence length (bp) | 1568 | 786 |
| N50 (bp) | 4919 | 1654 |

**Table S3 *In-cis* target genes of DE lncRNAs in S vs G locusts**

| **ID** | **Classification** | **Regulated** | **Target** | **Annotation** |
| --- | --- | --- | --- | --- |
| LNC473855.9 | antisense exonic | down | NA | NA |
| LNC582192.5 | antisense exonic | down | LOCMI09200 | NA |
| LNC115292.1 | antisense exonic | down | LOCMI17236 | small nuclear ribonucleoprotein sm D1 |
| LNC740388.9 | antisense exonic | down | LOCMI16374 | hypothetical protein SINV_01609 |
| LNC523317.1 | antisense exonic | down | LOCMI08000 | NA |
| LNC334351.1 | antisense exonic | down | NA | NA |
| LNC461737.1 | antisense exonic | down | NA | NA |
| LNC533775.2 | antisense exonic | down | NA | NA |
| LNC740388.2 | antisense exonic | down | LOCMI16374 | hypothetical protein SINV_01609 |
| LNC1364612.1 | antisense exonic | down | NA | NA |
| LNC302381.1 | antisense exonic | down | NA | NA |
| LNC823051.3 | antisense exonic | down | NA | NA |
| LNC712966.1 | antisense exonic | down | NA | NA |
| LNC740388.7 | antisense exonic | down | LOCMI16374 | hypothetical protein SINV_01609 |
| LNC292410.1 | antisense exonic | down | NA | NA |
| LNC794337.3 | antisense exonic | down | NA | NA |
| LNC1302682.1 | antisense exonic | down | LOCMI07182 | similar to AGAP000767-PA |
| LNC1296817.1 | antisense exonic | down | NA | NA |
| LNC359424.11 | antisense exonic | down | NA | NA |
| LNC1239462.10 | antisense exonic | down | LOCMI10539 | hypothetical protein KGM_15453 |
| LNC700550.1 | antisense exonic | down | NA | NA |
| LNC1186169.1 | antisense exonic | down | NA | NA |
| LNC1074886.1 | antisense intron | down | NA | NA |
| LNC444601.3 | antisense intron | down | LOCMI10088 | protein phosphatase 2C, putative |
| LNC1140096.6 | overlapping | down | NA | NA |
| LNC856134.1 | overlapping | down | NA | NA |
| LNC1274734.17 | overlapping | down | LOCMI17559 | NA |
| LNC387673.1 | overlapping | down | NA | NA |
| LNC1257221.1 | overlapping | down | NA | NA |
| LNC312400.1 | overlapping | down | NA | NA |
| LNC1330718.1 | overlapping | down | NA | NA |
| LNC522212.1 | overlapping | down | NA | NA |
| LNC721075.1 | overlapping | down | NA | NA |
| LNC448952.1 | overlapping | down | LOCMI06610 | NA |
| LNC1255087.2 | overlapping | down | LOCMI09290 | NA |
| LNC886555.1 | overlapping | down | NA | NA |
| LNC1151613.1 | overlapping | down | NA | NA |
| LNC168167.1 | overlapping | down | NA | NA |
| LNC160616.20 | sense exonic | down | LOCMI09558 | similar to GA18260-PA, partial |
| LNC386475.1 | sense exonic | down | NA | NA |
| LNC1033996.25 | sense exonic | down | NA | NA |
| LNC1033996.13 | sense exonic | down | NA | NA |
| LNC954646.1 | sense exonic | down | LOCMI09518 | NA |
| LNC1054722.3 | sense exonic | down | NA | NA |
| LNC988700.11 | sense exonic | down | LOCMI12633 | probable methyltransferase BCDIN3D-like |
| LNC1314926.11 | sense exonic | down | LOCMI16436 | isopentenyl pyrophosphate: dimethylallyl pyrophosphate isomerase |
| LNC1091246.117 | sense exonic | down | LOCMI08438 | NA |
| LNC1091246.113 | sense exonic | down | LOCMI08432 | NA |
| LNC1380899.9 | sense exonic | down | NA | NA |
| LNC1091251.1 | sense intron | down | LOCMI08430 | NA |
| LNC1384612.23 | sense intron | down | NA | NA |
| LNC463919.52 | sense intron | down | NA | NA |
| LNC481658.2 | sense intron | down | LOCMI10319 | similar to ankyrin 2,3/unc44 |
| LNC1263014.1 | sense intron | down | NA | NA |
| LNC1117914.1 | sense intron | down | LOCMI13121 | hypothetical protein KGM_21312 |
| LNC814165.2 | sense intron | down | NA | NA |
| LNC686212.30 | sense intron | down | NA | NA |
| LNC1409894.1 | sense intron | down | NA | NA |
| LNC428083.1 | sense intron | down | NA | NA |
| LNC1403702.1 | sense intron | down | NA | NA |
| LNC1390693.1 | sense intron | down | NA | NA |
| LNC150424.1 | sense intron | down | NA | NA |
| LNC1306651.1 | sense intron | down | NA | NA |
| LNC814165.5 | sense intron | up | NA | NA |
| LNC1201049.1 | sense intron | up | NA | NA |
| LNC1086477.2 | sense intron | up | NA | NA |
| LNC144811.24 | sense intron | up | NA | NA |
| LNC296820.13 | sense intron | up | NA | NA |
| LNC1286150.12 | sense intron | up | LOCMI07718 | similar to mCG1534 |
| LNC498536.1 | sense intron | up | LOCMI10339 | NA |
| LNC296820.7 | sense intron | up | NA | NA |
| LNC296820.10 | sense intron | up | NA | NA |
| LNC834441.36 | sense intron | up | NA | NA |
| LNC814165.4 | sense intron | up | NA | NA |
| LNC510762.6 | sense intron | up | NA | NA |
| LNC457607.2 | sense intron | up | NA | NA |
| LNC1210607.1 | sense intron | up | NA | NA |
| LNC1018051.107 | sense intron | up | LOCMI14877 | NA |
| LNC1255204.3 | sense intron | up | NA | NA |
| LNC1318798.4 | sense intron | up | NA | NA |
| LNC367075.1 | sense intron | up | NA | NA |
| LNC1151761.2 | sense intron | up | NA | NA |
| LNC1231026.15 | sense exonic | up | LOCMI04890 | NA |
| LNC344728.26 | sense exonic | up | NA | NA |
| LNC1166803.2 | sense exonic | up | NA | NA |
| LNC1159789.2 | sense exonic | up | LOCMI17261 | U2 small nuclear ribonucleoprotein A' |
| LNC283777.5 | sense exonic | up | NA | NA |
| LNC455821.14 | sense exonic | up | LOCMI06597 | coiled-coil domain-containing protein 16 |
| LNC1385449.1 | overlapping | up | NA | NA |
| LNC949134.1 | overlapping | up | NA | NA |
| LNC686137.5 | overlapping | up | LOCMI13560 | hypothetical protein BRAFLDRAFT_85516 |
| LNC1403702.2 | overlapping | up | NA | NA |
| LNC1018051.82 | overlapping | up | LOCMI14877 | NA |
| LNC192016.2 | antisense intron | up | NA | NA |
| LNC1251591.15 | antisense intron | up | LOCMI10740 | zinc finger protein 729-like |
| LNC342698.2 | antisense exonic | up | NA | NA |
| LNC965863.2 | antisense exonic | up | NA | NA |
| LNC286636.1 | antisense exonic | up | NA | NA |
| LNC986285.3 | antisense exonic | up | NA | NA |
| LNC311954.17 | antisense exonic | up | NA | NA |
| LNC1320041.1 | antisense exonic | up | LOCMI16291 | Hypothetical protein DAPPUDRAFT_302355 |

**Table S4** **Pearson correlation coefficient between early changed lncRNAs and all mRNAs in CS and IG**

See Supplementary Table 4.xlsx file.

**Table S5** **Correlations between known phase-related genes and lncRNAs**

| **Coding** | **LncRNA** | **r** | **p_value** | **Annotation** |
| --- | --- | --- | --- | --- |
| LOCMI02013 | LNC1065048.14 | 0.976838 | 0.00818 | *Vat1* isoform 2 |
| LOCMI03912 | LNC492755.1 | -0.95162 | 0.005784 | *Vat1* |
| LOCMI04527 | LNC1088763.7 | -0.97611 | 0.006934 | *NPF* |
| LOCMI16723 | LNC531328.2 | -0.9022 | 0.005805 | *NPYR* |
| LOCMI16723 | LNC1425451.2 | -0.90571 | 0.005428 | *NPYR* |
| LOCMI17298 | LNC494161.1 | -0.91876 | 0.00493 | *Ebony* |

**Table S6 Transcription factors co-expressed with LNC1010057**

| **Genes ID** | **Annotation** |
| --- | --- |
| LOCMI03997 | ZNF300 |
| LOCMI07647 | lolal |
| LOCMI08073 | Gtf2e2 |
| LOCMI08233 | ZNF677 |
| LOCMI09032 | brms1la |
| LOCMI09997 | ZNF808 |
| LOCMI10710 | Nfya |
| LOCMI13536 | TMF1 |

**Table S7 The neighboring genes of early changed lncRNAs**

| **LncRNA** | **Process** | ***In-cis* target gene** | **Annotation** | **Gene expression** |
| --- | --- | --- | --- | --- |
| LNC290551.1 | CS | LOCMI14917 | NA | No change |
| LNC750847.1 | CS | LOCMI10529 | NA | No change |
| LNC1309888.2 | CS | LOCMI15132 | farnesyl pyrophosphate synthase-like | No change |
| LNC515718.3 | CS | LOCMI05215 | similar to ENSANGP00000028549 | No change |
| LNC884242.64 | CS | LOCMI06585 | NA | No change |
| LNC918392.1 | CS | LOCMI04605 | NA | No change |
| LNC696144.1 | CS | LOCMI11517 | NA | No change |
| LNC492185.6 | CS | LOCMI10704 | RAB-28, putative | No change |
| LNC455821.12 | CS | LOCMI06598 | Charged multivesicular body protein 5 | No change |
| LNC270742.1 | CS | LOCMI05432 | Septin-4 | No change |
| LNC859600.5 | CS | LOCMI15272 | NA | No change |
| LNC1191349.1 | CS | LOCMI14085 | NA | No change |
| LNC481658.1 | CS | LOCMI10319 | similar to ankyrin 2,3/unc44 | No change |
| LNC532590.21 | CS | LOCMI07344 | protocadherin beta-11-like | No change |
| LNC1362214.3 | CS | LOCMI05222 | Cyclin-T | No change |
| LNC740388.3 | CS | LOCMI16374 | hypothetical protein SINV_01609 | No change |
| LNC409945.1 | CS | LOCMI09388 | NA | No change |
| LNC995027.1 | CS | LOCMI07055 | zinc finger protein, putative | No change |
| LNC1074881.1 | CS | LOCMI02608 | NA | No change |
| LNC582195.1 | CS | LOCMI09200 | NA | Change |
| LNC1159789.2 | CS | LOCMI17261 | Probable U2 small nuclear ribonucleoprotein A' | No change |
| LNC516660.6 | CS | LOCMI03916 | NA | No change |
| LNC1205086.1 | CS | LOCMI16463 | decapentaplegic protein | Change |
| LNC960219.6 | CS | LOCMI12404 | NA | No change |
| LNC965853.22 | CS | LOCMI06777 | NA | No change |
| LNC1065048.14 | CS | LOCMI14508 | guanine nucleotide-binding protein subunit beta-like | No change |
| LNC226217.1 | CS | LOCMI15512 | heat shock protein 20.7 | Change |
| LNC287252.1 | CS | LOCMI10563 | hypothetical protein KGM_15810 | No change |
| LNC494161.1 | CS | LOCMI09722 | hypothetical protein TcasGA2_TC015601 | No change |
| LNC222053.2 | CS | LOCMI17458 | odorant receptor 10 | No change |
| LNC488380.2 | CS | LOCMI04797 | uncharacterized protein C4orf14 homolog | No change |
| LNC1196508.7 | CS | LOCMI16450 | RAC protein kinase DRAC-PK85, putative | No change |
| LNC995231.4 | CS | LOCMI05033 | leucine-rich transmembrane protein, putative | No change |
| LNC427628.1 | CS | LOCMI11975 | Ankyrin repeat and sterile alpha motif domain-containing protein 1B | No change |
| LNC1018051.111 | CS | LOCMI14877 | NA | No change |
| LNC481658.2 | CS | LOCMI10319 | similar to ankyrin 2,3/unc44 | No change |
| LNC884242.43 | CS | LOCMI06585 | NA | No change |
| LNC521813.8 | IG | LOCMI07919 | mitochondrial manganese superoxide dismutase | No change |
| LNC1166374.22 | IG | LOCMI07685 | similar to CG7029 CG7029-PC | No change |
| LNC1362214.6 | IG | LOCMI05222 | Cyclin-T | No change |
| LNC494161.1 | IG | LOCMI09722 | hypothetical protein TcasGA2_TC015601 | No change |
| LNC913441.4 | IG | LOCMI04780 | hypothetical protein TcasGA2_TC003869 | No change |
| LNC752448.6 | IG | LOCMI04012 | serine/threonine-protein phosphatase 4 regulatory subunit 1 | No change |
| LNC1019444.4 | IG | LOCMI02750 | NA | No change |
| LNC1091251.1 | IG | LOCMI08430 | NA | No change |
| LNC1264076.19 | IG | LOCMI06937 | protein tipE-like | No change |
| LNC605645.1 | IG | LOCMI03761 | hypothetical protein SINV_16184 | Change |
| LNC1274734.17 | IG | LOCMI17559 | NA | No change |
| LNC183435.1 | IG | LOCMI09738 | NA | No change |
| LNC372428.3 | IG | LOCMI03063 | Arf-GAP with Rho-GAP domain, ANK repeat and PH domain-containing protein 2 | No change |
| LNC810363.1 | IG | LOCMI13311 | NA | No change |
| LNC732419.7 | IG | LOCMI08044 | NA | No change |
| LNC455821.4 | IG | LOCMI06597 | coiled-coil domain-containing protein 16 | No change |
| LNC115292.1 | IG | LOCMI17236 | small nuclear ribonucleoprotein sm D1, putative | No change |
| LNC867890.1 | IG | LOCMI12432 | hypothetical protein LOC100743128 | No change |
| LNC654627.19 | IG | LOCMI16951 | NA | No change |
| LNC1000175.2 | IG | LOCMI05004 | Phosphatidylinositide phosphatase SAC2 | No change |
| LNC1166430.1 | IG | LOCMI07690 | hypothetical protein KGM_09955 | No change |
| LNC1389123.3 | IG | LOCMI11374 | dual specificity protein phosphatase 12 | No change |
| LNC988700.11 | IG | LOCMI12633 | probable methyltransferase BCDIN3D-like | No change |
| LNC910465.2 | IG | LOCMI04470 | NA | No change |
| LNC1196508.8 | IG | LOCMI16450 | RAC protein kinase DRAC-PK85, putative | No change |
| LNC522395.8 | IG | LOCMI04472 | hypothetical protein SINV_03219 | No change |
| LNC177145.4 | IG | LOCMI03700 | NA | No change |
| LNC504062.1 | IG | LOCMI03584 | NA | No change |
| LNC1320041.1 | IG | LOCMI16291 | hypothetical protein DAPPUDRAFT_302355 | No change |
| LNC309610.1 | IG | LOCMI16041 | synaptic glycoprotein SC2, putative | No change |
| LNC322549.9 | IG | LOCMI05142 | E3 ubiquitin-protein ligase RFWD2 | No change |
| LNC989757.2 | IG | LOCMI14118 | Cytochrome c-type heme lyase | No change |
| LNC310207.1 | IG | LOCMI11753 | NA | No change |
| LNC1384975.3 | IG | LOCMI02446 | E3 ubiquitin-protein ligase RFWD2 | No change |
| LNC1364402.1 | IG | LOCMI07104 | MANF/CDNF-like protein | No change |
| LNC1013476.1 | IG | LOCMI15505 | heat shock protein 70 | No change |
| LNC468760.1 | IG | LOCMI11243 | NA | No change |
| LNC365152.1 | IG | LOCMI05045 | MLL1/MLL complex subunit KIAA1267-like | No change |
| LNC705616.1 | IG | LOCMI04449 | Magnesium-dependent phosphatase, putative | Change |
| LNC810651.1 | IG | LOCMI17197 | Protein mago nashi | No change |
| LNC994577.4 | IG | LOCMI03883 | clathrin coat assembly protein AP50 | No change |
| LNC1019444.5 | IG | LOCMI02749 | NA | No change |
| LNC1112140.3 | IG | LOCMI10054 | NA | No change |
| LNC437684.9 | IG | LOCMI06794 | similar to tubulin-specific chaperone e | No change |
| LNC972294.1 | IG | LOCMI16063 | fructose 1,6-bisphosphate aldolase | No change |
| LNC590493.64 | IG | LOCMI06800 | hypothetical protein LOC409034 | No change |
| LNC988705.1 | IG | LOCMI12632 | Beta-1,3-galactosyltransferase 5 | No change |
| LNC294462.1 | IG | LOCMI09497 | NA | No change |
| LNC673390.1 | IG | LOCMI14992 | NA | No change |
| LNC988759.1 | IG | LOCMI12634 | NA | No change |
| LNC960219.3 | IG | LOCMI12405 | NA | No change |
| LNC794623.1 | IG | LOCMI04068 | hypothetical protein | No change |
| LNC498536.1 | IG | LOCMI10339 | NA | No change |
| LNC1009861.1 | IG | LOCMI11558 | jerky protein homolog-like | No change |
| LNC358214.1 | IG | LOCMI16893 | gustatory receptor 45 | No change |
| LNC1264089.58 | IG | LOCMI06937 | protein tipE-like | No change |
| LNC1137408.25 | IG | LOCMI08581 | NA | Change |
| LNC1143850.107 | IG | LOCMI13993 | NA | No change |
| LNC929723.1 | IG | LOCMI10569 | similar to AGAP005618-PA | No change |

**Table S8 Primers used in this study**

| Primer name | Sequence, 5′-3′ | Description |
| --- | --- | --- |
| LNC1010057R1 | CTCGCAACCCATCACACAAC | 5′ RACE |
| LNC1010057R2 | GGACCTCACAGTTCCAACAC | 5′ NEST RACE |
| LNC1010057F1 | GTGTGGCGTTTTCTTGTAGGG | 3′ RACE |
| LNC1010057F2 | CCCCTTGCCAAAGAATGGAAG | 3′ NEST RACE |
| LNC992414R1 | CCATCTTTCAATATGCTACACACCC | 5′ RACE |
| LNC992414F1 | GATAGGGTTCACATCCATCCCG | 3′ RACE |
| LNC992414F2 | CCAAGCCCACTATAAGGACTTACC | 3′ NEST RACE |
| Repeat-RNAi-F1-365 | ATGGGACGAGATAGGCTGTAAGAG | RNAi  RNAi  RNAi  RNAi  RNAi  RNAi  RNAi  RNAi |
| Repeat-RNAi-R1-552 | CCATTACTAACCACCAGTCA |  |
| Repeat-RNAi-F1-521 | ACAGAGTATGATGACTGGTGG |  |
| Repeat-RNAi-R1-945 | CATATACTACTAAGGGGATC |  |
| LNC992414-RNAi-F1 | TCTTTGGGAGGTGTAGCAATT |  |
| LNC992414-RNAi-F2 | CAACCTGACCACACTCATTCG |  |
| LNC992414-RNAi-R1 | AATGAGTGTGGTCAGGTTGAA |  |
| LNC992414-RNAi-R2 | CAATATGGTTGCTCCTGGTCT |  |
| Repeat-RT-F1-142 | AATGTCGTTGAGTGGAAGG | qRT-PCR  qRT-PCR  qRT-PCR  qRT-PCR  qRT-PCR  qRT-PCR  qRT-PCR  qRT-PCR  qRT-PCR  qRT-PCR  qRT-PCR  qRT-PCR  qRT-PCR  qRT-PCR  qRT-PCR  qRT-PCR  qRT-PCR  qRT-PCR  qRT-PCR  qRT-PCR |
| Repeat-RT-R1-552 | CCATTACTAACCACCAGTCA |  |
| LNC992414-RT-F | GGTTGAACAGAGTCTCAGAGT |  |
| LNC992414-RT-R | ACTGTGGGATAGTTGGGTCAT |  |
| LNC756712F | ACAGTGAGGTCCCTATGAAATG |  |
| LNC756712R | CCAAACTAACCAGGTCATAC |  |
| LNC992415F | ATTTGGCTACAATAACACGC |  |
| LNC992415R | CCGCAGGAATTGATACCTACATAAG |  |
| LNC324366F | AGTTGAGGTTCATACCGAAG |  |
| LNC324366R | CAATGCTGCCCTTCTTTCTG |  |
| LNC1365633F | ATTTCTGAACACGGTACAAG |  |
| LNC1365633R | GTAAAGACGCCCGCTCAGTTG |  |
| LNC1379668F | TTACTGTGCCTGGGAAACCAAC |  |
| LNC1379668R | TGACATGGCTGACTGAGAAG |  |
| LNC531294F | GGTCTTCGCACTTATTCAAC |  |
| LNC531294R | TCAGACTTCGCCTTAACAGG |  |
| LNC643475F | CTTTCTACCGTCTGTCTTCG |  |
| LNC643475R | TGGAAATGGTGCCAAACGAC |  |
| LNC775093F | TGGGCAAGTGAGTTAGTACATC |  |
| LNC775093R | TACCACTCTCACAAGCAATCT |  |
